# Spatial and temporal control of expression with light-gated LOV-LexA

**DOI:** 10.1101/2021.10.19.465021

**Authors:** Inês M.A. Ribeiro, Wolfgang Eßbauer, Romina Kutlesa, Alexander Borst

**Affiliations:** Max Planck Institute of Neurobiology Am Klopfersptiz 18, 82152 Martinsried, Germany

## Abstract

The ability to drive expression of exogenous genes in different tissues and cell types, under control of specific enhancers, has been crucial for discovery in biology. While many enhancers drive expression broadly, several genetic tricks were developed to obtain access to isolated cell types. Studies of spatially organized neuropiles in the central nervous system of insects have raised the need for a system that targets subsets of cells within a single neuron type, a feat currently dependent on stochastic flip-out methods. To access the same subsets of cells within a given expression pattern consistently across fruit flies, we developed the light-gated expression system LOV-LexA. We combined the bacterial LexA transcription factor with the plant-derived light oxygen voltage (LOV) photosensitive domain and a fluorescent protein. Exposure to blue light uncages a nuclear localizing signal in the C-terminal of the LOV domain, and leads to translocation of LOV-LexA to the nucleus, with subsequent initiation of transcription. LOV-LexA enables spatial and temporal control of expression of transgenes under LexAop sequences in larval fat body as well as pupal and adult neurons with blue light. The LOV-LexA tool is ready to use with GAL4 and Split-GAL4 drivers in its current form, and constitutes another layer of intersectional genetics, that provides light-controlled genetic access to specific subsets of cells across flies.

## Introduction

Patterned expression of genes is essential for differentiation of distinct cell types during development. Enhancers defining expression patterns have long been used in binary expression systems to study development and function of specific cell types. Binary expression systems couple enhancer-led expression of an exogenous transcription factor to expression of a transgene that sits downstream of promoter sequences exclusively bound by the exogenous transcription factor (DEL VALLE Rodriguez *et al.* 2011). The GAL4-UAS system uses the yeast transcription factor GAL4 under control of an enhancer, that binds upstream activating sequences (UAS), which in turn drive expression of transgenes sitting downstream UAS (Brand and Perrimon 1993). Random insertions of P-elements carrying GAL4 into the genome were used to trap enhancers, with expression of GAL4 dependent on neighboring regions in the genome (Rubin and Spradling 1982; Spradling and Rubin 1982; Bellen *et al.* 1989; Grossniklaus *et al.* 1989; Wilson *et al.* 1989; Perrimon *et al.* 1991; Venken and Bellen 2012; Venken and Bellen 2014). More recently, stretches of noncoding genomic DNA carved out of known gene enhancers, or from regions predicted to contain enhancers, have been extensively used to generate large collections of driver lines (Pfeiffer *et al.* 2008; Pfeiffer *et al.* 2010; Jenett *et al.* 2012; Kvon *et al.* 2014; YANEZ-Cuna *et al.* 2014; Tirian and Dickson 2017). Other binary expression systems were added to the fruit fly genetic toolbox. The LexA-LexAop (Lai and Lee 2006) and the QF-QUAS (Potter *et al.* 2010; Riabinina *et al.* 2015) systems both rely on exogenous transcription factors and DNA binding sequences, and can be combined with GAL4-UAS, allowing for independent access to multiple cell types in the same organism (e.g. Feng *et al.* 2014; Sen *et al.* 2017; Ribeiro *et al.* 2018; Feng *et al.* 2020). The spatial resolution, or cell type-specificity of binary expression systems, is determined by the enhancer driving expression of the exogenous transcription factor. Given that it is still not possible to design enhancers specific for many cell types (Serebreni and Stark 2021), it is necessary to screen to obtain enhancers specific for the cell type of interest.

Several methods, under the umbrella of intersectional genetics, were developed to further restrict transgene expression in binary systems. The modular nature of the GAL4 activation and DNA-binding domains enables separation of GAL4 into two parts, with each split-GAL4 half placed under the control of a different enhancer (Luan *et al.* 2006). The final transgene expression occurs only in cells that express both split-GAL4 halves, which dimerize through added leucine zipper domains to form a fully functional transcription factor. Existing collections of split-GAL4 lines targeting single neuron types were established by screening for enhancer pairs that together provide exclusive access to specific cell types of interest (e.g. Wu *et al.* 2016; Dionne *et al.* 2018; Namiki *et al.* 2018; Dolan *et al.* 2019; Schretter *et al.* 2020; Wang *et al.* 2020; Sterne *et al.* 2021). Other powerful methods of restricting expression to single or fewer cells include the recombinase-based systems for stochastic labeling (Lis *et al.* 1983; Golic and Lindquist 1989; Xu and Rubin 1993; Lee and Luo 1999; Hadjieconomou *et al.* 2011; Nern *et al.* 2015;

ISAACMAN-Beck *et al.* 2020), temperature sensitive mutations of Gal80, the repressor of GAL4 (Nogi *et al.* 1977; Lee and Luo 2001; Mcguire *et al.* 2003), and use of transcription factors modified to drive transcription in the presence of an ingestible drug (Mcguire *et al.* 2004). In addition to providing temporal and spatial control however, these methods either involve increase in temperature, that unleashes a stress response in all cells of the organism (LINDQUIST 1986), or addition of drugs with potential off-target effects, both of which may affect experimental outcomes.

With several recent advances in optics and laser technology, light is now easily modulated at the level of its spectrum, intensity and even beam shape (Chen *et al.* 2018). The high spatial and temporal precision of light pulse delivery to living organisms has the potential to take the spatial and temporal resolution of transgene expression to new levels. Several photosensitive proteins have been introduced into exogenous expression systems to amass the advantages of light as a precise trigger (DI Ventura and Kuhlman 2016; DE Mena *et al.* 2018; DI Pietro *et al.* 2021).

Phytochromes (Phy) are sensitive to red and far-red light, and bind the phytochrome interacting factor (PIF) in presence of light (Yamamoto and Deng 1999). The Photo-GAL4 tool capitalizes on the PhyB and PIF light-dependent interaction to reconstitute a complete GAL4 upon exposure to light (DE MENA AND RINCON-LIMAS 2020). To function, Photo-GAL4 requires addition of phycocyanobilin (PCB), a chromophore that is absent in animal cells (Yamamoto and Deng 1999), limiting its applicability (DE MENA AND RINCON-LIMAS 2020). The cryptochrome split-LexA system similarly uses cryptochrome 2 (CRY2) and its binding partner, the cryptochrome interacting protein (CIB), to gate reformation of split-LexA with blue light (Szuts and Bienz 2000; Chan *et al.* 2015). In ShineGal4, the pMagnet and nMagnet photoswitches derived from the blue light photoreceptor VVD endogenous to *Neurospora crassa*, replace the leucine zippers in split-GAL4 halves and heterodimerize upon exposure to light (Kawano *et al.* 2015; DI Pietro *et al.* 2021). ShineGal4 functions in several epithelia across developmental stages, with its current form limited to a few drivers.

To circumvent these limitations and expand the photosensitive toolbox in *Drosophila*, we developed a light-gated expression system based on the light, oxygen or voltage (LOV) domain originally found in oat phototropin 1 (*Avena sativa*) (Christie *et al.* 1998; Christie *et al.* 1999; Crosson and Moffat 2002) and LexA (Horii *et al.* 1981; WALKER 1985; Rhee *et al.* 2000; Masuyama *et al.* 2012), under the control of UAS sequences. LOV-LexA gates expression of transgenes with blue light *in vivo*, in several cell types in larval, pupal, and adult fruit flies. LOV-LexA can be directly crossed to split-GAL4 and GAL4 drivers, adding thus another layer of spatiotemporal control to transgene expression in *Drosophila* that is combinable with existent binary expression systems, and transferable to other model organisms.

## Materials and Methods

### Plasmids and cloning

The LexA chimeras LexA:GAD, LexA:p65 and LexA:VP16 from the plasmids pBPLexA::GADUw, pBPLexA::p65Uw and pBPLexA::VP16Uw (Addgene # 26230, 26231, 26232, Gerald Rubin lab) were mutagenized to change the NLS-like sequence from: (2433) GTT ACT GTG AAA CGT CTC AAG AAG CAA GGC AAT (VTVKRLKKQGN), to: (2433) GTT ACT GTG AAA <u>G</u>GG CTC <u>G</u>AG AAG CAA GGC AAT (VTVK<u>G</u>L<u>E</u>KQGN) (Rhee *et al.* 2000), using the Q5 site directed mutagenesis kit (New England Biolabs, catalogue # E0554S). The resulting modified LexA (mLexA) chimeras were combined through DNA assembly (Gibson *et al.* 2009) with the following components: eLOV (Addgene # 92213, Alice Ting lab) (Wang *et al.* 2017), SV40 nuclear localizing signal (Pfeiffer *et al.* 2010) and tdTomato (Shaner *et al.* 2004) or GFP (from pJFRC7-20XUAS-IVS-mCD8::GFP , Addgene # 26220, Gerald Rubin lab) (Pfeiffer *et al.* 2008) or FLAG (amino acid sequence: DYKDDDDK) with a kit (Gibson assembly kit from New England Biolabs, catalogue # E5510S). The different combinations were cloned into pJFRC7-20XUAS-IVS-mCD8::GFP (Addgene # 26220) cut with XhoI (NEB catalogue # R0146S) and XbaI (NEB catalogue # R0145S), to replace mCD8::GFP, and produce pJFRC7- 20XUAS-LexA-transactivator-eLOV-tag construct combinations.

### S2R+ cell culture, transfection, stimulation, fixation, and immunostaining

The *Drosophila* cell line S2R+ (ECHALIER 1997) was obtained from the Drosophila Genomics Resource Center, supported by NIH grant 2P40OD010949. S2R+ cells were cultured at 25°C in Schneider’s Medium (Gibco, cat # 21720-024) containing 10% fetal bovine serum (Gibco, cat # A47668-01) and 1% penicillin-streptomycin (Gibco, cat # 15070-063). To test the various LexA- transactivator-eLOV-tag constructs, listed in supplemental Table S1, for cell survival and ability to drive expression gated by light, S2R+ cells were transfected with pMET-GAL4 as the driver, the UAS-LexA-transactivator-eLOV-tag test construct, and pJFRC19-13XLexAop2-IVS-myr::GFP (Addgene #26224) or 13XLexAop-IVS-myr::tdTomato (this study) as the LexAop-led reporters of LexA-transactivator-eLOV-tag transcriptional activity. We used co-transfection of pMET-GAL4 (Velichkova *et al.* 2010), pJFRC7-20XUAS-IVS-mCD8::GFP (Pfeiffer *et al.* 2008), pJFRC7-20XUAS-IVS-mCherry (this study) as controls to characterize transfection efficiency of three constructs simultaneously. Three DNA plasmids, 200 to 250ng/µl, were combined with FuGene (Promega, cat # E2311) in Schneider’s media with a proportion of 600 to 750 ng DNA for 4ul FuGene. The DNA plasmid/FuGene mix was allowed to stand for 30 minutes to one hour at room temperature, after which it was added to roughly 1 million cells pre-plated in a 24-well plate. The metallothionein promoter in the pMET-GAL4 driver (Velichkova *et al.* 2010) is activated by addition of copper sulfate (CuSO4, Sigma-Aldrich Nr. 451657), to a final concentration of 0.75 mM. Presentation of light was initiated 1 to 3 hours after addition of copper sulfate. Light was delivered in pulses of 30s of blue LED (from the LED light source of the inverted laboratory microscope LEICA DM IL LED) at 1 Hz (Table S3). Cells were fixed with 4% paraformaldehyde overnight at 4°C, between 8 to 10 hours after addition of copper sulfate, and processed for immunostaining with standard protocols (e.g. Velichkova *et al.* 2010), with antibodies anti-GFP chicken antibody dilution 1:2000 (Rockland, catalogue # 600-901-215S; RRID: AB_1537403), anti-RFP rabbit antibody dilution 1:2000 (Rockland, catalogue # 600-401- 379, RRID: AB_11182807), anti-FLAG rat antibody dilution 1:300 (Novus Biologicals, catalogue # NBP1-06712, RRID: AB_1625981). Secondary antibodies were goat anti-chicken Alexa Fluor 488 1:1000 (Thermo Fischer Scientific catalogue # A-11039; RRID: AB_2534096), goat anti-rabbit Alexa Fluor 568 1:1000 (Thermo Fischer Scientific catalogue # A-11011; RRID: AB_143157, and goat anti-rat Alexa Fluor 568 1:1000 (Thermo Fischer Scientific catalogue # A- 11077; RRID: AB_2534121).

### Quantification of signal intensity in cell culture

Five images per well, in 24-well plates, were obtained from immuno-stained cells with an inverted fluorescence microscope (Leica DM IL LED), with a 5x objective, with green and red filter cubes. The open-source software CellProfiler (version 4.1.3) (Carpenter *et al.* 2006; Mcquin *et al.* 2018) was used to segment individual cells in each image, based on the test construct fluorescent tag signal with the Otsu method, and measure the amount of LexAop-reporter in each segmented cell, as a proxy for transcription levels of LexAop reporters by test constructs, LexA- transactivator-eLOV-tag. We used the mean intensity of segmented cells in Cell Profiler, Object MeanIntensity, as a measure of mean pixel intensity per segmented cell, in the green and red channels. This measure is a normalized value by default in Cell Profiler and is plotted in figure panels 1C, 1D, S1B and S1D. Scripts written in Python (version 3.8, http://www.python.org) were then used to read data values per Object, average all segmented cells across at least two independent experiments and plot the data.

### *Drosophila* culture and genetics

Fly stocks obtained from the Bloomington Drosophila Stock Center (NIH P40OD018537) were used in this study. All the strains of the fruit fly *Drosophila melanogaster* used in this study are listed in supplemental Table S2. Fruit flies were maintained on standard cornmeal-agar medium supplemented with baker’s yeast and incubated at 18°C or 25°C with 60% humidity and 12h light/dark cycle. Males and females were tested indiscriminately throughout experiments. Larvae of the second and third instar were used for tests on fat body with *Cg-GAL4*; 2 to 4 days after puparium formation (APF) pupae were used for tests in neurons in the central brain and dorsal abdominal oenocytes; adults ranging from 1 to 6 days old were used for tests in adult neurons.

### Light Stimulation in fat body

To test UAS-LOV-LexA, UAS-eLOV-nls-tdTomato-mLexA:GAD and UAS-eLOV-nls-tdTomato-mLexA:VP16 constructs in fat body cells, second to third instar larvae from crosses with *Cg-GAL4* (Asha *et al.* 2003), were removed from the food, washed in water and placed in a well with 40 µl of 15% sucrose in water solution, one larva per well in a 96-well plate wrapped in aluminum foil to shield the larvae from light. Pulses of blue light were delivered to individual wells, with the 96-well plate mounted on an inverted microscope. Light from a blue LED (LEICA DM IL LED) was delivered at 11.7mW, at 1Hz for 30s (Table S3). Half the plate was not exposed to light and larvae in such wells served as controls. The fat bodies were dissected 6 to 12 hours after light delivery, fixed and immunostained with anti-GFP chicken antibody dilution 1:1000 (Rockland, catalogue # 600-901-215S; RRID: AB_1537403) and anti-RFP rabbit antibody dilution 1:2000 (Rockland, catalogue # 600-401-379, RRID: AB_11182807). Secondary antibodies were goat anti-chicken Alexa Fluor 488 1:1000 (Thermo Fischer Scientific catalogue # A-11039; RRID: AB_2534096), goat anti-rabbit Alexa Fluor 568 1:1000 (Thermo Fischer Scientific catalogue # A-11011; RRID: AB_143157). Stained fat bodies were mounted with Vectashield Antifade Mounting Medium (BIOZOL, Ref H-1000) and imaged with a Leica TCS SP8 confocal microscope.

### Light Stimulation in neurons

To stimulate neurons in pupal stages, pupae were recovered from vials by adding water to the vial wall to dissolve the glue binding pupal cases to the pupation site. Pupae aged between 2 and 3 days after pupal formation (APF) were then dried on a kimwipe tissue and glued on a double-side sticky tape spread on a cover slip (Figure 3C), that was attached to a microscope slide with plasticine. Adult flies were glued to a custom-made aluminum or plastic plate with a hole, with a diameter ranging from 300 to 400 µm, large enough to expose part of the head of the adult fly and shield the rest of the fly from light. Melted Eicosane 99% (Aldrich 219274-5G) was added to the thorax and part of the head to immobilize the adult fly and shield part of the head from light (Figure 4F). Such custom holders were mounted on a microscope slide with plasticine, separating the flies from the slide.

Slides bearing pupae or adult flies were then mounted on an upright confocal microscope (Leica TCS SP8) for pre-programmed serial light delivery with the 458nm laser at 10% power at 5.75µW (Table S3). Each light pulse was composed to 30 to 50 scans across a depth of 200 to 400 µm. Using the xyzt mode of Leica software together with position mapping, it was possible to deliver light serially to many pupae or adult flies prepped together. After light delivery, cover slips with pupae were vertically inserted into new food vials and incubated for 2 days for expression in neurons in the fly brain, before dissection. Adult flies were removed from holders after light delivery, by breaking the brittle Eicosane, placed in fresh food vials and incubated at 25°C for 1 to 2 days before dissection.

Adult brains were dissected in cold PBS, fixed in 4% PFA for 20 to 40 minutes at room temperature, washed in PBS with 0.5% Triton X-100 two times, incubated with DAPI (Invitrogen, D1306) at dilution 1:3000 in PBS with 0.5% Triton X-100 for 10 minutes, washed again, mounted with Vectashield Antifade Mounting Medium (BIOZOL, Ref H-1000), and imaged on the same day with a Leica TCS SP8 confocal microscope.

### Live imaging in oenocytes and neurons

The pupal case covering the most anterior abdomen in the case of oenocytes (*w; 109(2)-GAL4, UAS-CD8:GFP/+; UAS-LOV-LexA/+*), or the head in case of neurons (*w, LexAop-CsChrimson:Venus;+;UAS-LOV-LexA/fru-GAL4)*, of pupae lined up on a double side sticky tape on a slide (see above, Figure 4A), was removed under low light conditions, or as low as possible since pupal cuticle is transparent. The slide was mounted on a Leica TCS SP8 confocal microscope, and positions for serial imaging were marked. The pupae were then kept in the dark for 30 minutes before live imaging was initiated.

Oenocytes were firstly scanned in the red channel alone, followed by exposure to blue light (see Table S3). Afterwards, oenocytes were imaged in the red channel every five minutes for at least one hour and 40 minutes. Oenocytes were imaged one last time in the red and green channels, obtain CD8:GFP signal to delineate the oenocyte cell body, in addition to LOV-LexA. To image *fru+* neurons in pharate adult pupae (4 d APF), a scan in the green and red channels preceded the exposure to light (see Table S3), after which pupal heads were scanned in the red and green channels every hour. Most pupae eclosed after 12 to 14 hours under the confocal microscope.

### Quantification of signal intensity in flies

Mounted fat body and brain tissue were imaged using a Leica TCS SP8 confocal microscope with a 20.0X objective, using the lasers 405, 488 and 568 to image DAPI, Venus and tdTomato respectively. The same laser power, gain and line averaging were used within each experiment in order to compare fluorescence levels across light and dark conditions. Z-stacks thus obtained were cropped in XY and Z with ImageJ/Fiji (Schindelin *et al.* 2012), to isolate cell bodies expressing the test construct. Z-projections of these crops were loaded with the Scikit-image image processing package into Python (version 3.8, http://www.python.org), to obtain pixel intensity values in a 2- dimensional matrix for each color channel in RGB. The ratio of the mean pixel intensity in green and red channels, or green and blue channels, was used to compare relative fluorescence levels of reporter gene to test constructs.

## Results

### Design of an expression system gated by light

The LOV2 domain of *Avena sativa* phototropin 1, AsLOV2, is photosensitive (Harper *et al.* 2003; Lungu *et al.* 2012; Zayner *et al.* 2012; Diensthuber *et al.* 2014). Exposure to blue light causes the Jɑ helix to unfold, thereby freeing its C-terminus (Harper *et al.* 2003). This property arises from interactions with flavin, the blue light-absorbing chromophore present in animal cells, and can be used to expose a small peptide of up to 10 amino-acid residues long, added to or integrated into the Jɑ C-terminus (Huala *et al.* 1997; Christie *et al.* 1998; Christie *et al.* 1999; Salomon *et al.* 2000; Harper *et al.* 2003). This photosensitive system has been used to cage several peptides in genetic tools, including the nuclear localizing signal (NLS) to shuttle proteins to the nucleus, the tobacco etch virus protease (TEVp) cleavage site for an integrator of neuronal activity and reporters of protein-protein interactions (Wang *et al.* 2010; Strickland *et al.* 2012; MOTTA- Mena *et al.* 2014; Niopek *et al.* 2014; Guntas *et al.* 2015; Yumerefendi *et al.* 2015; Jayaraman *et al.* 2016; Niopek *et al.* 2016; Yumerefendi *et al.* 2016; Reade *et al.* 2017; Smart *et al.* 2017; Salinas *et al.* 2018; VAN Haren *et al.* 2018; Zhao *et al.* 2018; Cavanaugh *et al.* 2020). Recent work employed directed evolution on the native AsLOV2 to develop the evolved LOV (eLOV), that presents improved stability in the dark state due to three single nucleotide mutations (Kim *et al.* 2017a; Wang *et al.* 2017; Kim *et al.* 2019). We added the short NLS from SV40 (Pfeiffer *et al.* 2010), to make eLOV-nls, and regulate availability of the NLS to the cell milieu with blue light (Niopek *et al.* 2014).

To build a transcription factor gated by light, we selected the binary expression system LexA/LexAop (Szuts and Bienz 2000; Loewer *et al.* 2004; Lai and Lee 2006), that is complementary to the widespread GAL4-UAS system and has been successfully incorporated in diverse model organisms (Lai and Lee 2006; Emelyanov and Parinov 2008; NONET 2020).

LexA is a repressor of transcription endogenous to *Escherichia coli* (Horii *et al.* 1981), where it regulates the SOS response (WALKER 1985). Addition of an activation domain to the C-terminal of LexA renders such LexA-transactivator chimeras capable of activating transcription of transgenes sitting downstream of the LexA operator (LexAop) (Rhee *et al.* 2000; Lai and Lee 2006). In *Drosophila*, the most common LexA-transactivator chimeras contain the activation domains GAL4 activation domain (GAD, LexA:GAD), p65 (LexA:p65) or VP16 (LexA:VP16) (Rhee *et al.* 2000; Szuts and Bienz 2000; Lai and Lee 2006; Emelyanov and Parinov 2008; Yagi *et al.* 2010). Despite its bacterial origin, LexA carries an NLS-like sequence that allows it to shuttle to the nucleus when expressed in eukaryotic cells (Rhee *et al.* 2000; Pfeiffer *et al.* 2010; Masuyama *et al.* 2012). To make translocation of LexA to the nucleus solely dependent on eLOV- nls, we mutagenized the NLS-like sequence in the LexA codon optimized for *Drosophila melanogaster* (Rhee *et al.* 2000), and created a modified LexA (mLexA) (Figure 1A, see Materials and Methods). We examined the propensity to translocate to the nucleus of mLexA-transactivator chimeras by transfecting such constructs into the S2R+ *Drosophila* cell line (ECHALIER 1997), together with the metallotheionein-GAL4 (Met-GAL4) that drives ubiquitous expression upon addition of CuSO4 (Velichkova *et al.* 2010), and the reporter myr:GFP under control of LexAop sequences (Pfeiffer *et al.* 2010) (Figure 1B). All three chimeras of unmodified LexA- transactivator drove expression of the LexAop-myr:GFP to levels similar to UAS-CD8:GFP (Figure 1C), confirming their ability to shuttle to the nucleus (Rhee *et al.* 2000; Pfeiffer *et al.* 2010; Masuyama *et al.* 2012). In contrast, mLexA-transactivator chimeras led to reduced expression of the reporter transgene (Figure 1C), confirming that the NLS-like sequence in LexA plays a major role in shuttling LexA to the nucleus.

**Figure 1:**
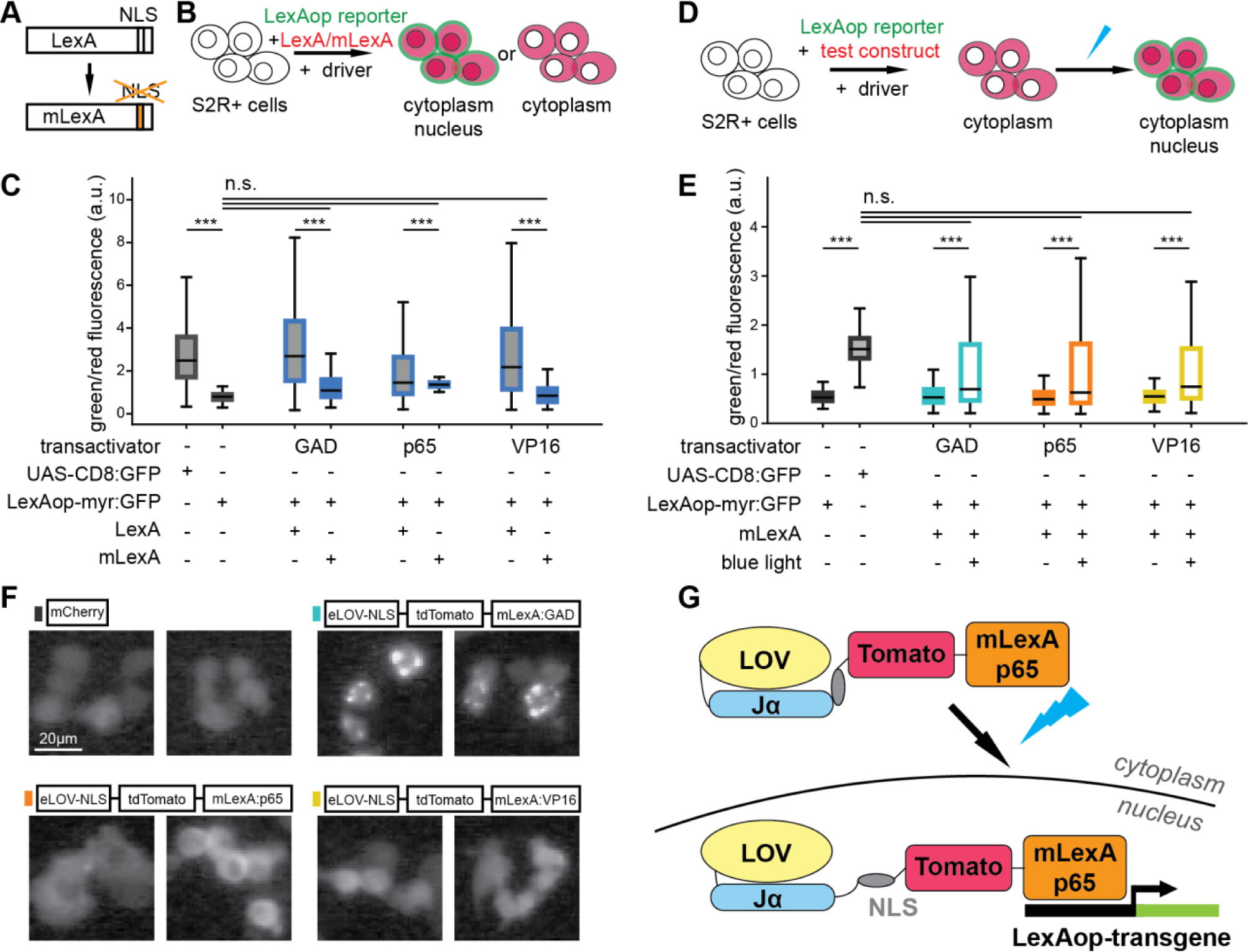
Testing components for a light-gated expression system based on eLOV. **A.** The NLS-like sequence in LexA is mutagenized in modified LexA (mLexA). **B.** S2R+ cell line was used to test whether LexA-transactivator:tdTomato and mLexA:tdTomato drive transcription of a LexAop-reporter, LexAop-myr:GFP, using the pMET-GAL4 driver. **C.** The ratio of LexAop reporter myr:GFP expression in relation to expression of the different LexA- and mLexA- transactivator chimeras determined by tdTomato fluorescence. Co-transfection of UAS-mCherry and UAS-CD8:GFP was used as an approximate measure of co-expression (first bar in the boxplot), whereas co-transfection of UAS-mCherry with the LexAop reporter LexAop-myr:GFP established the baseline (second bar in the boxplot). The constructs for GAD were LexA:GAD- tdTomato and mLexA:GAD-tdTomato, for p65 were LexA:p65-tdTomato and mLexA:p65-tdTomato, and for VP16 were LexA:VP16-tdTomato and mLexA:VP16-tdTomato. Mutagenizing NLS-like sequence reduces transcriptional activity of mLexA-transactivator chimeras compared to LexA-transactivator chimeras. At least 200 cells with medium levels of expression of mCherry or tdTomato, from at least two transfections of S2R+ cells are represented for each condition. **D.** S2R+ cells were transfected with a reporter, LexAop-myr:GFP, together with the driver pMET- GAL4 and the test construct, to examine light-gated transcription for test constructs. **E.** Expression of LexAop reporter myr:GFP in relation to expression of mLexA-transactivator chimeras combined with eLOV. Placement of eLOV-nls N-terminal followed by the fluorescent protein tdTomato and the mLexA-transactivator chimera yielded the best signal for cells exposed to pulses of blue light, while maintaining a low reporter signal in cells kept in the dark. As in B, co- transfection of UAS-mCherry with UAS-CD8:GFP (second bar in plot) or LexAop-myr:GFP (first bar in plot) served as a positive and negative control, respectively. The test constructs were eLOV- nls-tdTomato-mLexA:GAD (GAD), eLOV-nls-tdTomato-mLexA:p65 (p65) and eLOV-nls- tdTomato-mLexA:VP16 (VP16). At least 200 cells with medium levels of expression of mCherry or tdTomato, from 2 to 5 transfections of S2R+ cells are represented for each condition. **C, E.** *** represents p values < 0.001, n.s. represents p values > 0.05, obtained with Student’s t test. **F.** Representative examples of S2R+ cells expressing test constructs indicated above the images under control of pMET-GAL4. The eLOV-nls-tdTomato-mLexA:GAD forms clusters in the cytoplasm, whereas both eLOV-nls-tdTomato-mLexA:p65 and eLOV-nls-tdTomato- mLexA:VP16 are evenly distributed in the cytoplasm, and sometimes nucleoplasm, like mCherry. **G.** Schematic showing how eLOV-nls-tdTomato-mLexA:p65 (or LOV-LexA) works.

We combined eLOV-nls with the three mLexA-transactivator chimeras, and a fluorescent protein (Shaner *et al.* 2004), placed each combination under control of UAS and tested their performance in S2R+ cells co-transfected with Met-GAL4 and LexAop-myr:GFP for constructs tagged with a red fluorescent protein or LexAop-myr:tdTomato for constructs tagged with GFP (Figure S1A,C). Several of the mLexA constructs carrying eLOV-nls at the C-terminal led to expression of myr:GFP in the dark (Figure 1E, Supplemental Figure S1B,D), indicating that NLS is frequently uncaged with eLOV-nls at the C-terminal end, even in the absence of light. On the other hand, many of the mLexA constructs with eLOV-nls N-terminal were unable to drive expression of myr:GFP upon presentation of blue light (Supplemental Figure S1B,D). The combinations made with LexA:GAD chimera formed clusters in the cytoplasm irrespective of the fluorescent protein used as a tag (Supplemental Figure S1E,H), while most other combinations were homogenously distributed in the cytoplasm and occasionally in the nucleoplasm (Figure 1F, and Supplemental Figure S1F-J). Of note, cells with high levels of expression of many of the mLexA constructs tested, presented expression of myr:GFP irrespective of the light regime delivered (data not shown), indicating that eLOV is unstable if expressed at high levels, as previously observed (Kim *et al.* 2017a; Kim *et al.* 2019). Cells expressing eLOV-nls-tdTomato-mLexA:p65 and eLOV-nls-tdTomato-mLexA:VP16 at moderate levels, presented no to very little LexAop-myr:GFP reporter expression in the dark, and displayed increase in expression of LexAop-myr:GFP upon exposure to blue light (Figure 1E). These two constructs thus gathered the characteristics necessary for a light-gated expression system and were used to create transgenic flies. Despite its shortcomings, the eLOV-nls-tdTomato-mLexA:GAD was also injected since mLexA:GAD is suppressible by Gal80, potentially providing another level of regulation of a light-gated expression system.

### Characterization of eLOV-nls-tag-mLexA chimera constructs *in vivo*

*Drosophila* larvae have transparent cuticle that allows for internal tissues to be exposed to unabated light. The bilateral, multilobed fat body running along the larva, is visible underneath the body wall musculature, and is targeted by the collagenase enhancer (Cg-)GAL4 (Asha *et al.* 2003). Distribution of eLOV-nls-tdTomato-mLexA:GAD, eLOV-nls-tdTomato-mLexA:p65 or eLOV-nls-tdTomato-mLexA:VP16 in larval fat body followed the trend observed in S2R+ cells (Figure 2A,D, and Supplemental Figure S2A-F), with eLOV-nls-tdTomato-mLexA:GAD forming clusters (Supplemental Figure S2C,D) and eLOV-nls-tdTomato-mLexA:p65 or eLOV-nls-tdTomato-mLexA:VP16 distributing evenly in the cytoplasm, and occasionally in the nucleoplasm (Figures 2C,D,F,G, and Supplemental Figure S2F,G). To test the ability to induce expression of a reporter under control of LexAop sequences, LexAop-CsChrimson:Venus (Klapoetke *et al.* 2014) (hereafter referred to as Venus), second and third instar larvae reared at 18°C were placed in 96 well plates in a 15% sucrose solution, to repress their tendency to wander, and exposed to several pulses of low intensity blue light (Figure 2A,B, supplemental Table S3). Larvae were then incubated at 25°C for 7 to 11 hours, after which the fat body was dissected, fixed, and stained. Despite considerable expression of eLOV-nls-tdTomato-mLexA:GAD and eLOV-nls-tdTomato- mLexA:VP16 in fat body cells, exposure to blue light failed to elicit expression of the reporter (Supplemental Figure S2A-H). In contrast, exposure of larvae expressing eLOV-nls-tdTomato- mLexA:p65 to as little as 3 pulses of blue light, led to increase in reporter expression under control of LexAop sequences (Figure 2A-I). Surprisingly, a lower number of pulses of blue light combined with longer incubation at 25°C resulted in maximum increase in reporter expression (Figure 2I). This suggests that despite low intensity, exposure to too many pulses of blue light leads to less efficiency of light-gated expression. Given that expression of eLOV-nls-tdTomato-mLexA:p65 in S2R+ and fat body cells kept in the dark presented no or very low expression of the reporter gene and that exposure to blue light led to increase in reporter expression, the construct eLOV-nls-tdTomato-mLexA:p65 was selected for further studies and named LOV-LexA (Figure 1G).

**Figure 2.**
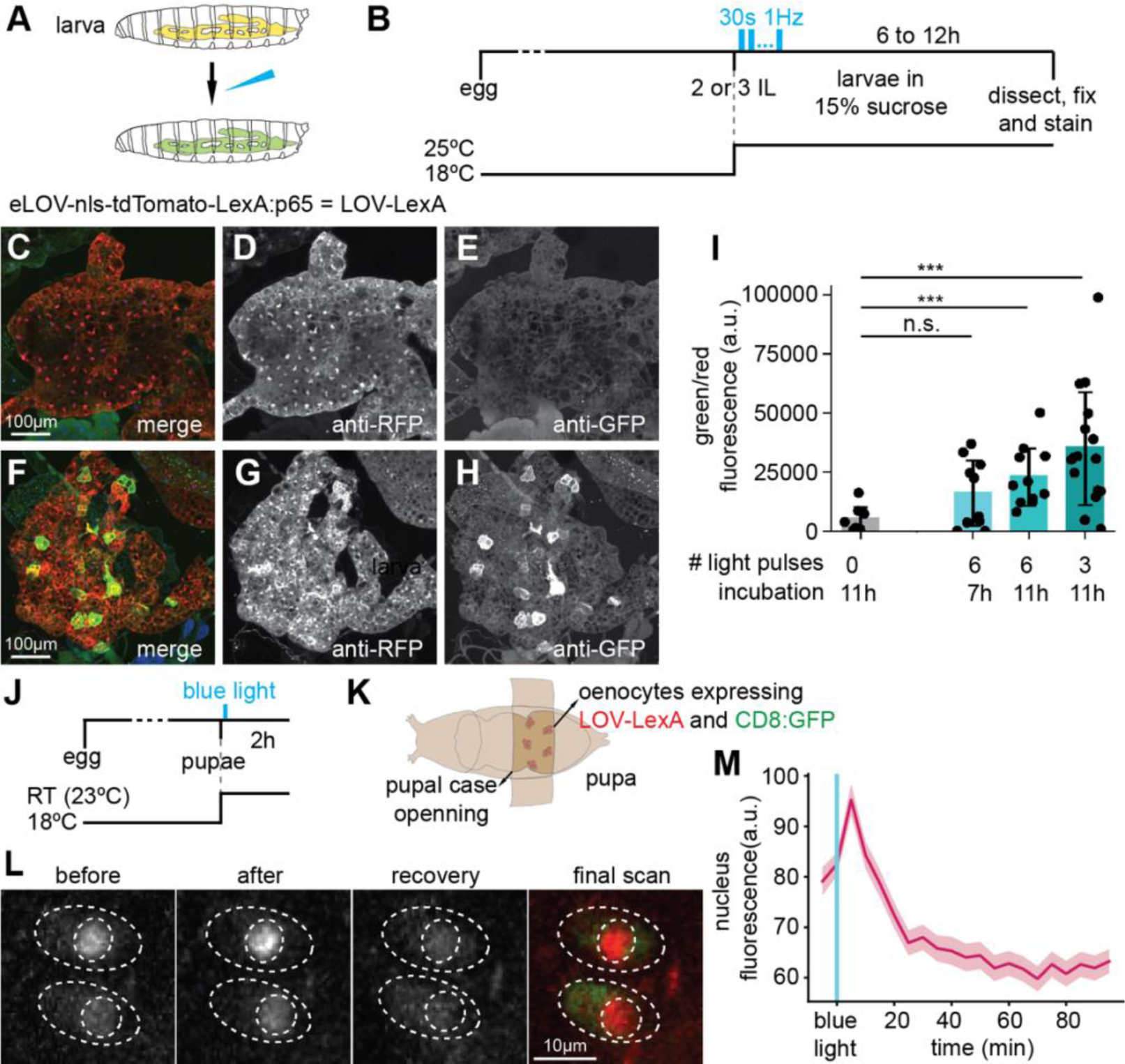
LOV-LexA is gated by light *in vivo*. **A.** *Drosophila* larvae expressing LOV-LexA in the fat body were exposed to blue light and examined for expression of the LexAop reporter, as well as the construct selected for LOV-LexA. **B.** Schematics showing the timeline of the experiment, light regime. Second or young third instar larvae were selected from vials kept at 18°C and transferred to 15% sucrose solution. The first light pulse was delivered immediately after this transfer, with a blue LED on an inverted microscope (supplemental Table S3). Larvae were placed in the dark at 25°C between light pulses (see text for details), until dissection. **C-H.** Fat bodies expressing LOV-LexA with Cg-GAL4 for second to third instar larvae kept in the dark (C-E) or exposed to three 30s pulses of blue LED light at 1 Hz (F-H). Exposure to blue light appears to alter LOV-LexA cellular distribution (D, G) and leads to expression of LexAop-CsChrimson:Venus in fat body cells as detected with anti-GFP antibody (E, H). **I.** Ratio of fluorescence, measured as pixel intensity in confocal-acquired images, of anti-GFP signal/anti-RFP signal for stained fat bodies from larvae with the genotype *w, LexAop- CsChrimson:Venus; Cg-GAL4/+; UAS-LOV-LexA/+* that were kept in the dark (N=7, representative example in C-E), exposed to 6 light pulses and dissected after 7 hours (N=10), or 11 hours (N=11), or exposed to 3 light pulses and dissected 11 hours later (N=16, representative example in F-H). Varying the number of light pulses and the incubation period at 25°C before dissection led us to conclude that LOV-LexA gates expression with light in fat body, and that LOV-LexA light-gated expression is highest with three light pulses and an 11-hour incubation period at 25°C. *** represents p-values < 0.001, n.s. represents p-values > 0.05, two-tailed Mann-Whitney tests. Exposure to blue light leads to an increase in the amount of Venus relative to LOV- LexA levels. **J.** Schematics showing the timeline of the experiment, light exposure, and functional imaging. **K.** *Drosophila* pupae expressing LOV-LexA and CD8:GFP in oenocytes were mounted on double-side sticky tape, and an opening in the pupal case that exposes oenocytes, was created. Pupae expressed LOV-LexA and CD8:GFP in oenocytes with the following genotype: *w; 109(2)- GAL4, UAS-CD8:GFP/+; UAS-LOV-LexA/+*. **L.** Representative images of pupal oenocytes showing LOV-LexA before (before) and immediately following exposure to blue light (after), 60 min after exposure to blue light (recovery) and 120 min after light exposure (final scan). The final scan included the green channel to capture CD8:GFP, co-expressed with LOV-LexA, and used to delineate the cell body. The light used to capture GFP is blue and elicited another translocation of LOV-LexA to the nucleus, thereby demonstrating that the oenocytes were healthy after imaging. **M.** Mean nuclear tdTomato fluorescence over time, imaged live every 5 min. Shades represent standard error of the mean (s.e.m.). LOV-LexA translocates to the nucleus upon exposure to blue light within minutes in oenocytes, and slowly leaks out of the nucleus after exposure to blue light.

The use of the AsLOV2 domain to cage a NLS signal has been previously demonstrated to effectively move coupled proteins into the nucleus in a light-dependent manner (Niopek *et al.* 2014; Yumerefendi *et al.* 2015). To determine the kinetics of LOV-LexA nuclear translocation, we expressed LOV-LexA in oenocytes, which are large cells sitting underneath the cuticle with roles in secretion and metabolism (Makki *et al.* 2014). Adult oenocytes arise in pupae and reach their final locations through several bouts of migration during metamorphosis. We imaged stationary oenocytes in pupae aged between 2 to 3 days after pupal formation (APF), through the transparent cuticle, after removal of the overlying pupal case (Figure 2J,K). After preparation of the samples, LOV-LexA was present in the cytoplasm as well as in the nucleus in abdominal oenocytes (Figure 2L ‘before’). Exposure to blue light (485nm, 2.53 µW, 30 slices, supplemental Table S3) leads to a rapid accumulation of LOV-LexA in the nucleus, that decreases over time (Figure L ‘after’, ‘recovery’, M). To determine the location of the cytoplasm, cells were imaged to detect CD8:GFP as well as tdTomato in LOV-LexA 100 minutes after exposure to blue light (Figure 2L ‘final scan’, not depicted in the graph in Figure 2M). LOV-LexA accumulated again in the nucleus in all oenocytes imaged (n=14), indicating that the reduction of LOV-LexA levels in the nucleus over time is not due to general degradation of the live preparation. LOV-LexA thus exhibits fast translocation to the nucleus, which peaks 5 minutes after exposure to blue light, and a slower movement out of the nucleus, reaching minimum levels after 20 minutes in the dark.

### LOV-LexA behavior in diverse neuronal types

Similar to fat body, we assessed LOV-LexA behavior in neurons with the transgene Venus under control of LexAop sequences (LexAop-CsChrimson:Venus) (Klapoetke *et al.* 2014) as a reporter of LOV-LexA transcriptional activity. Presence of CsChrimson:Venus is readily detected in its native expression in neurons with fixation alone, thereby eliminating the need for the extra amplification step of antibody immunostaining (Mckellar *et al.* 2019). We tested LOV-LexA in the lobula columnar 10 (LC10)-group neurons, LC10a, b, c, and d, that arborize in the lobula and project to anterior optic tubercle, in the dorsal fly brain (Otsuna and Ito 2006; Costa *et al.* 2016; Panser *et al.* 2016; Wu *et al.* 2016). LC10a neurons, but not LC10b, c, or d, mediate tracking of visual objects (Ribeiro *et al.* 2018; HINDMARSH Sten *et al.* 2021). Expression of LOV-LexA in LC10-group neurons with *LC10s-SS2* and *LC10a-SS1* drivers (Ribeiro *et al.* 2018) led to moderate expression of Venus in the dark if flies were raised at 25°C (Supplemental Figure S3A-E), but not if flies were raised at 18°C in the dark (Supplemental Figure S3F-J). This indicates that the dark state of LOV-LexA is unstable in flies reared at 25°C. The leakiness of LOV-LexA at 25°C could arise from an elevated accessibility to the NLS at higher temperatures, or increased LOV-LexA expression as previously observed in S2R+ cells (Supplemental Figure S1) and in other eLOV- based tools (Kim *et al.* 2017a; Kim *et al.* 2019). The stability of LOV-LexA in the dark was further tested with the panneuronal driver *GMR57C10-GAL4* (Jenett *et al.* 2012). In many neuron types, rearing flies at 18°C prevented accumulation of the Venus reporter in flies expressing LOV-LexA panneuronally (Supplemental Figure S3K-M). Several neuron types, including neurons in the optic lobe, mushroom body, antennal lobe and suboesophageal region, were an exception to this rule and presented high levels of Venus expression. Differences in expression strength across neuron types represented in the *GMR57C10-GAL4* expression pattern partially account for the observed variability in Venus expression in the dark. On the other hand, differential expression pattern of genes involved in nucleocytoplasmic transport in different neuron types could potentially underlie these discrepancies. Alpha importins function as adaptors that bind NLS peptides, bringing proteins with NLS in contact with β importins which in turn, mediate transport into the nucleus. The ɑ importin *ɑ Karyopherin 4* (*ɑKap4*, CG10478) is highly expressed in Kenyon cells and other neuron types (Supplemental Figure S3N) (Venken *et al.* 2011; Larkin *et al.* 2021). Expression of the ɑ importin karyopherin ɑ1 (Kap- ɑ1, CG8548) and the ß importins cadmus (cdm, CG7212) and Chromosome segregation 1 (Cse1, CG13281) are limited to a small number of neuron types in the central brain (Supplemental Figure S3O-Q). The presence of *ɑKap4*, or other importins, in certain neuron types could potentially explain the selected leakiness of LOV-LexA dark state. To test this, we co-expressed *Kap-ɑ1* (Jang *et al.* 2015; Larkin *et al.* 2021) with LOV-LexA in LC10a-SS1 neurons in flies reared at 18°C in the dark. Co-expression of LOV-LexA with Kap-ɑ1 in LC10a-SS1 neurons led to expression of Venus reporter gene (Supplemental Figure S3R-V), suggesting that increase in nucleocytoplasmic transport may facilitate translocation of LOV-LexA to the nucleus, in the dark.

We tested several GAL4 and split-GAL4 drivers in flies raised at 18°C and compared native expression of LOV-LexA and the reporter Venus. Like in other cell types, above certain levels of expression of LOV-LexA, the amount of Venus detected in neurons correlated with that of LOV- LexA (Supplemental Figure S3Z). Together these observations suggest that the LOV-LexA tool has a stable dark state in drivers of weak to moderate expression strength, which constitute the majority of GAL4 and split-GAL4 lines available for genetic access to single neuron types.

### LOV-LexA mediates light-gated expression in neurons

The pupal case and the adult cuticle are tanned and block light, leading to decreased exposure of internal tissues to light. To uncage the NLS in LOV-LexA expressed in pupal and adult brain, we used a 1-photon laser with 458 nm wavelength in a confocal microscope (see Materials and Methods and Table S3). The driver *fru-GAL4*, a *GAL4* knock-in in the locus of the gene *fruitless* (*fru*) (Gailey and Hall 1989; Stockinger *et al.* 2005), targets approximately 100 neuron types, collectively called *fru* neurons, many of which were shown to regulate courtship behavior (among others, Cachero *et al.* 2010; Yu *et al.* 2010; Lu *et al.* 2012; Thistle *et al.* 2012; Toda *et al.* 2012; Bath *et al.* 2014; Inagaki *et al.* 2014; Ribeiro *et al.* 2018; Mckellar *et al.* 2019).

Expression is initiated in pupal development with low expression strength at late pupal stages. Pupae reared at 18°C and expressing LOV-LexA in *fru* neurons, were exposed to a series of four pre-programmed light pulses (supplemental Table S3), after which they were placed at 25°C for two days (Figure 3A-C). Expression of Venus was significantly increased in most *fru* neurons in pupae that were exposed to blue light (Figure 3D-H). Importantly, pupae kept in the dark displayed little or no expression of Venus (Figure 3D,E). Similar outcomes were observed for the *LC10a- SS1* driver. Like *fru-GAL4*, *LC10a-SS1* drives expression in pupal stages at low levels (Ribeiro *et al.* 2018, and data not shown). Exposure of pupae to four pulses of 1-photon laser 458nm light spaced over 30 minutes (supplemental Table S3), elicited light-dependent expression of Venus in LC10a neurons (Figure 3I-M). Delivery of four to eight pulses of blue light, but not two, proved to be sufficient for appreciable increase in Venus expression (Figure 3N). Increase in expression of the reporter Venus in fru+ and LC10-group neurons exposed to light, and its absence in the same neurons kept in the dark, demonstrates that LOV-LexA gates expression with blue light in neurons in the pupal brain.

**Figure 3:**
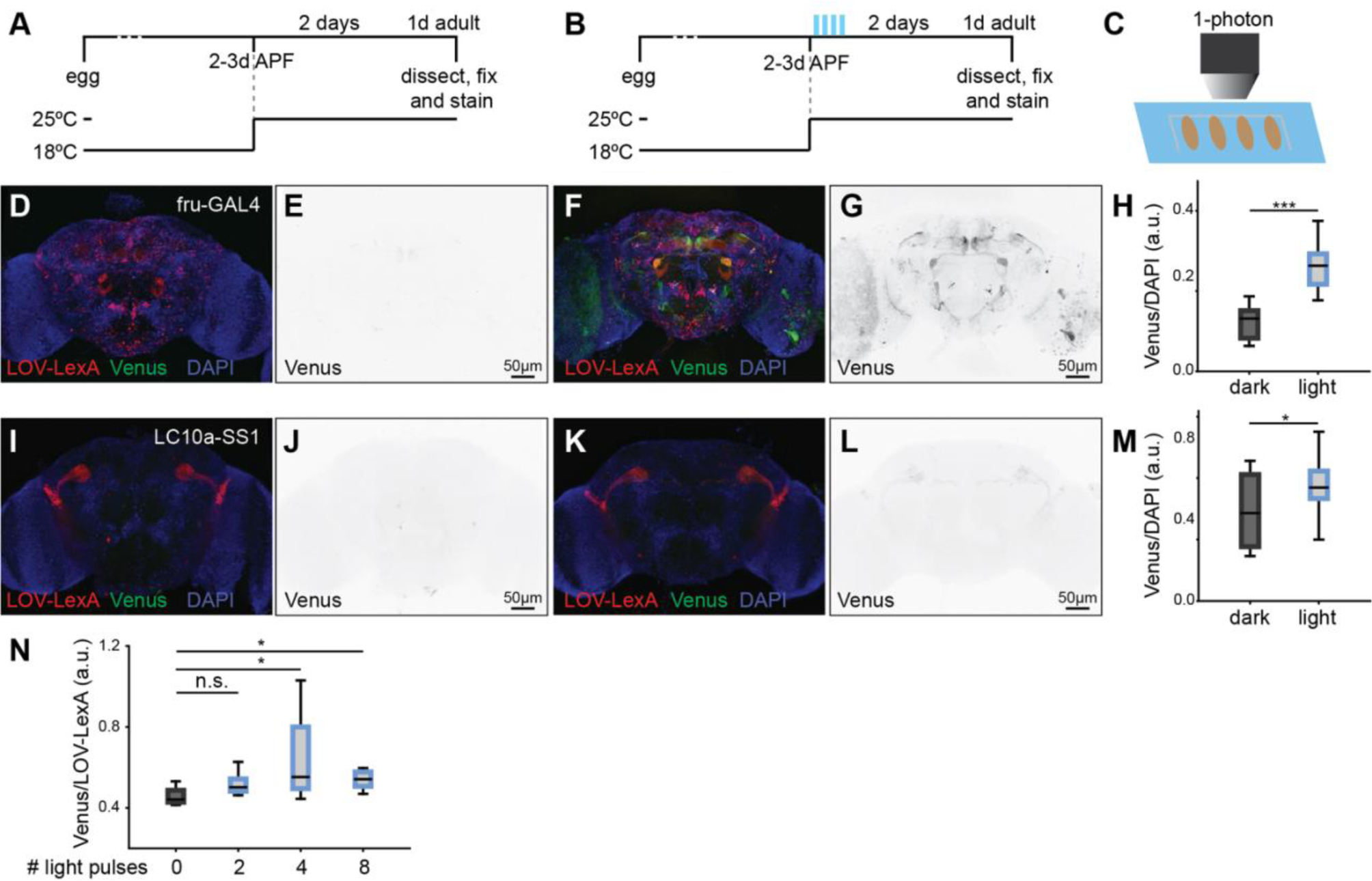
LOV-LexA gates expression with light in neurons. **A, B.** Schematic outlining the experiment. Pupae reared at 18°C aged 2 to 3d APF were removed from vials, mounted on double side sticky tape on a cover slip and kept in the dark (A) or pasted onto a slide and exposed to blue light (B). Mounted pupae kept in the dark or exposed to light were shifted to 25°C until dissection. **C.** Schematic showing pupae lined on double side sticky tape for light delivery. **D - G.** Adult brains showing native expression of LOV-LexA (red in D and F) and LexAop-CsChrimson:Venus (Venus, green in D and F, and dedicated image in E and G) from pupae kept in the dark (D, E) or exposed to light (F,G) at 3-4 days APF, as shown in B. **H.** Ratio of Venus signal intensity over DAPI signal intensity for *fru+* neuronal cell bodies located in the anterior brain. Pupae exposed to pulses of blue light (N=12) express the LexAop reporter Venus at higher levels compared to pupae kept in the dark (N=13), demonstrating that exposure to light leads to higher LOV-LexA transcriptional activity. **I-L.** Adult brains showing native expression of LOV-LexA (red in I and K) and LexAop-CsChrimson:Venus (Venus, green in I and K, and dedicated image in J and L) from *w, LexAop-CsChrimson:Venus;+/LC10a-SS1.AD;UAS-LOV- LexA/LC10a-SS1.DBD* pupae kept in the dark (I, J) or exposed to light at 3-4 days APF (K, L), as shown in B. **M.** Ratio of Venus signal intensity over DAPI signal intensity for LC10a neuronal cell bodies. Pupae exposed to pulses of blue light (N=12) express the LexAop reporter Venus at higher levels compared to pupae kept in the dark (N=6). **N.** Ratio of Venus over LOV-LexA native fluorescence from adult brains *w, LexAop-CsChrimson:Venus;+;UAS-LOV-LexA/fru-GAL4* exposed to 0, 2, 4 or 8 pulses of blue light as 2 to 3 d APF pupae (N=3, 4, 11, 5 respectively). *** represents p-values < 0.001, * represents p-values < 0.05, n.s. represents p-values > 0.05, two- tailed Mann-Whitney tests.

Precise control of the time of initiation of transgene expression has numerous advantages, including allowing for embryonic and pupal development to occur undisturbed in the absence of ectopic expression and for regulation of the level of transgene expressed. We measured the time it takes for LOV-LexA to drive transcription of LexAop controlled Venus after exposure to blue light. The head in pupae expressing LOV-LexA with *fru-GAL4* was uncovered by removing the encapsulating pupal case and exposed to pulses of blue light (supplemental Table S3, Figure 4A). The pupal brain was then imaged every hour for 12 hours to determine the timing at which Venus starts to be expressed. Venus expression doubled 12 hours after blue light pulse delivery (Figure 4B-D). Detection of expression with native protein fluorescence in adult brains was reliably observed 24 hours after exposure to blue light during late pupal stages (Figure 3D-H), indicating that LOV-LexA light-gated expression takes 12 to 24 hours to accumulate enough LexAop Venus reporter to be visualized with native levels.

**Figure 4:**
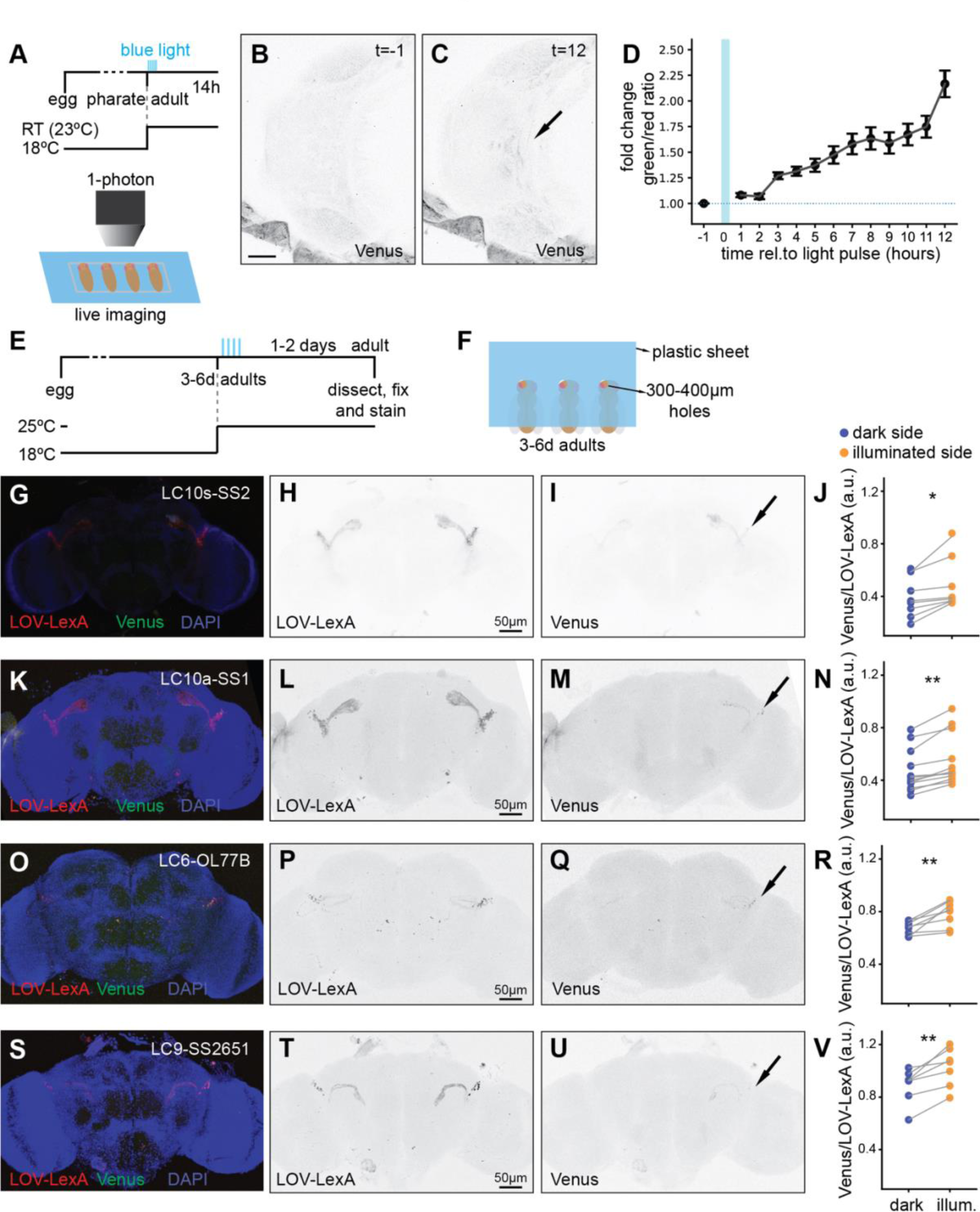
LOV-LexA enables spatial and temporal control of transgene expression with light. **A.** Schematic outlining the experiment (top) and schematic showing pupae lined up on a slide, with exposed heads for live imaging and blue light delivery (bottom), shown in C and D. **B,C.** Live image of a 4d APF pupal head after removal of the pupal case, with expression of Venus in fru+ neurons (*w, LexAop-CsChrimson:Venus;+;UAS-LOV-LexA/fru-GAL4*) before delivery of blue light (B) and 12 hours after delivery of blue light (C). **D.** Change in the ratio of native Venus signal over LOV-LexA tdTomato native signal, before and after light delivery (N=5). **E.** Timeline of the experiment. **F.** Schematic showing preparation to deliver spatially restricted light to immobilized adult flies, glued with low temperature melting wax to an opaque coverslip, with the head placed under a hole with a diameter between 300 to 400µm, shown in E to Q. **E-V.** Representative images of adult brains expressing LOV-LexA in several LC neurons and spatially restricted LexAop-CsChrimson:Venus, after exposure to spatially restricted blue light to target visual projection neurons unilaterally and quantification. **G-J.** LC10-group neurons LC10s-SS2 (*w, LexAop-CsChrimson:Venus; +/LC10s-SS2.AD; UAS-LOV-LexA/LC10s-SS2.DBD*) with N=8. **K-N.** LC10a neurons LC10a-SS1 (*w, LexAop-CsChrimson:Venus; +/LC10a-SS1.AD; UAS-LOV- LexA/LC10a-SS1.DBD*) with N=11. **O-R.** LC6-OL77B neurons (*w, LexAop-CsChrimson:Venus; +/OL77B.AD; UAS-LOV-LexA/OL77B.DBD*) with N=8. **S-V.** LC9-SS2651 neurons (*w, LexAop-CsChrimson:Venus; +/SS2651.AD; UAS-LOV-LexA/SS2651.DBD*) with N=7. **J,N,R,V.** Ratio of native Venus over native LOV-LexA (tdTomato) signals between the side of the head that was illuminated compared to the side that was kept in the dark, with each plot corresponding to the genotypes shown in the same row. * represents p-values < 0.05, ** represents p-values < 0.01, Wilcoxon test.

Drivers that initiate expression at adult stages, such as LC10s-SS2, were exposed to light at adult stages. Adult flies expressing LOV-LexA in LC10-group neurons were immobilized with low-melting wax on a custom-made opaque plastic coverslip with 300 to 400µm holes (Figure 4E,F). The area of cuticle above the cells of interest was placed under one of the holes (Figure 4F). Somata for the LC10-group neurons are located in the dorso-posterior side of the head, in an area bordering the rim of the retina. Immobilized flies with the cuticle covering somata of LC10-group neurons on one side of the adult head exposed, were delivered four to six pulses of 485nm 1-photon laser light over the course of one hour (supplemental Table S3). Detection of native fluorescence revealed accumulation of Venus in LC10-group neurons exclusively on the side exposed to light (Figure 4G-J). Similar light deliveries to adult flies expressing LOV-LexA in LC10a (Figure 4K- N), LC6 (Figure 4O-R) and LC9 neurons (Figure 4S-V) resulted in unilateral Venus expression. Importantly, most flies prepared in this fashion showed unilateral expression in LC neurons (Figure 4J,N,R,V), indicating that LOV-LexA allows for consistent genetic access to the same subsets of cells within an expression pattern.

## Discussion

We developed LOV-LexA, a light-gated expression system based on the photosensitive eLOV domain (Wang *et al.* 2017), and the modified transcription factor mLexA (Emelyanov and Parinov 2008; Pfeiffer *et al.* 2010). In the absence of light, LOV-LexA proteins reside in the cytoplasm of larval and adult cells. Delivery of blue light causes the LOV Jɑ helix to uncage an NLS, which then mediates translocation of LOV-LexA to the nucleus. Once in the nucleus, LOV- LexA drives expression of transgenes under control of LexAop sequences. The use of light as a trigger enables control of expression with high spatial and temporal resolution in live larvae and adult flies, making LOV-LexA an important addition to the *Drosophila* genetic toolbox that will expand the use of existent broad drivers as well as allow targeting subsets of cells within single cell types.

Several forms of LexA-transactivator chimeras are used in different animal models (Lai and Lee 2006; Emelyanov and Parinov 2008; NONET 2020). Surprisingly, the ability to remain outside the nucleus in the dark and to elicit reporter expression upon light exposure varied widely among different combinations of mLexA-transactivator chimeras, eLOV-nls and fluorescent tag. Replacing tdTomato with the FLAG tag in LOV-LexA, to make eLOV-nls-FLAG-mLexA:p65, leads to high levels of leakiness in the dark in S2R+ cells (data not shown), suggesting that intra-protein interactions among the different components of LOV-LexA play an important role in stability of the Jɑ helix in the dark (Kim *et al.* 2017a; Wang *et al.* 2017). Experiments in cell culture suggest that at high levels of expression, LOV-LexA proteins are more likely to translocate to the nucleus and drive expression of the LexAop reporter transgene. Rearing flies expressing LOV-LexA at 25°C similarly leads to unwanted expression of the LexAop reporter, imposing limits on the temperature used to raise fruit flies and the available driver lines. Further improvements of the eLOV domain have to be implemented to circumvent this limitation (Kim *et al.* 2019). Roughly 12 to 24 hours separate delivery of blue light and accumulation of LexAop transgene expression in neurons, giving the fly time to recover from potential adverse effects of delivery of blue light, that include temporary blindness (MONTELL 2012). This temporal separation might preclude use of transgenes encoding proteins with a short half-life. However, this time allows for other light and genetic manipulations to be performed on the same animal, without the need to perform all manipulations simultaneously on a tethered fly (Kim *et al.* 2017b).

Replacing the transcription factor in LOV-LexA with QF2 (Riabinina and Potter 2016), testing NLS sequences of varied strengths, or using other LOV-based domains might improve stability of LOV-LexA at higher temperatures and expression levels, and change the time required for reporter expression. Addition of another protein domain that counterbalances nuclear import, such as a nuclear export signal (Niopek *et al.* 2014) or a membrane tethering domain (Kim *et al.* 2017a), might provide more stability to LOV-LexA. On the other hand, some neuron types present LexAop reporter expression even if LOV-LexA is expressed at low levels. The uneven expression of importins across neuron types in the fly brain, similar to what is observed in the mouse brain (Hosokawa *et al.* 2008), suggests that different neuron types might express nucleocytoplasmic transport machinery to different extents. We predict that this variability is likely to influence how LOV-LexA functions across neuron and cell types, making cells with high nucleocytoplasmic transport capabilities less suitable for light-gated expression with LOV-LexA.

Other light-gated expression systems have been developed in *Drosophila*, including the cryptochrome split-LexA, Photo-Gal4 and ShineGal4 (Chan *et al.* 2015; DE MENA AND RINCON-LIMAS 2020; DI Pietro *et al.* 2021). We expressed the cryptochrome split-LexA with the same driver used to test LOV-LexA, LC10s-SS2 (Ribeiro *et al.* 2018), and found that cryptochrome split-LexA system is leaky in flies raised at 18°C and kept in the dark (Supplemental Figure S3X,Y). Given that Photo-GAL4 relies on PhyB and requires addition of the chromophore PCB, normally absent in animal cells, it is currently limited to *ex vivo* studies (DE MENA AND RINCON-LIMAS 2020). The chromophore providing LOV with light sensitivity, flavin, exists in animal cells, making the LOV-LexA system solely dependent on delivery of light. The limited number of enhancers driving ShineGal4, mostly targeting embryonic and pupal epithelia, prevents its widespread testing without re-cloning under other promoters. LOV-LexA is currently under control of the UAS promoter, and is one cross away from being tested with the myriad of GAL4 and split-GAL4 driver lines available.

There are thousands of enhancer-LexA or -GAL4 drivers targeting several cell types simultaneously (Jenett *et al.* 2012; Kockel *et al.* 2016; Robie *et al.* 2017; Tirian and Dickson 2017; Kockel *et al.* 2019). The LOV-LexA can be placed downstream of broadly expressed enhancers, in order to restrict transgene expression in the cell type of interest. Moreover, LOV- LexA downstream of an enhancer can be combined with GAL4 and QF binary expression systems, to genetically target two or more single neuron types independently in the same animal, enabling several different experiments, including simultaneous monitoring of neuronal activity or determining dependency relationships among different neuron types. Many neuron types are composed of dozens of cells that are topographically organized to represent the visual field (Fischbach and Dittrich 1989; Otsuna and Ito 2006; Wu *et al.* 2016). Topographic organization of neuropiles processing sensory information is also observed in other animals, like the mouse superior colliculus, visual cortex, and for other sensory modalities, like the barrel cortex (Huberman *et al.* 2008; Nassi and Callaway 2009; PETERSEN 2019), among others. LOV-LexA is an ideal tool to test the role of topography, by providing consistent genetic access to the same subsets of somata within a single neuron type, with little stochasticity. We demonstrate consistent targeting of several LC neurons unilaterally with LOV-LexA by targeting their somata. Applying this strategy to all visual projection neurons will elucidate how each contributes to guiding visual behavior.

Compared to *Drosophila melanogaster*, many model organisms in which it is possible to create transgenics, have smaller repertoires of enhancer driver lines that give access to different tissues and cell types. Implementing LOV-LexA in such model organisms will greatly amplify the number of specific cell types that can be genetically manipulated, expanding the landscape of possible experiments in emerging model organisms and the knowledge we can acquire from them.

### Tool and data availability

The DNA plasmid for LOV-LexA is deposited in DGRC (stock # 1583) and is available upon request. The *Drosophila melanogaster* LOV-LexA flies will be deposited in VDRC (stock # 311200), and are also available upon request. Data sets are available upon request. Please contact Inês M.A. Ribeiro at.

## Acknowledgements

We are grateful to M.Sauter and C.Theile for technical assistance with fly husbandry, S.Prech for assistance with sets ups used to deliver light, R.Kasper and E.Laurell for managing the Imaging Facility, to A.Fabritius for assistance with cloning, L.Groschner for assistance with mounting adult flies, and to A.H.Ali, R.M.Vieira and G.Ammer for critically reading the manuscript. The Bloomington Drosophila Stock Center, the *Drosophila* Genomics Resource Center, VDRC and FlyBase were instrumental for this research project and countless others, and the authors wish these institutions remain fully funded.

---

**Figure S1:**
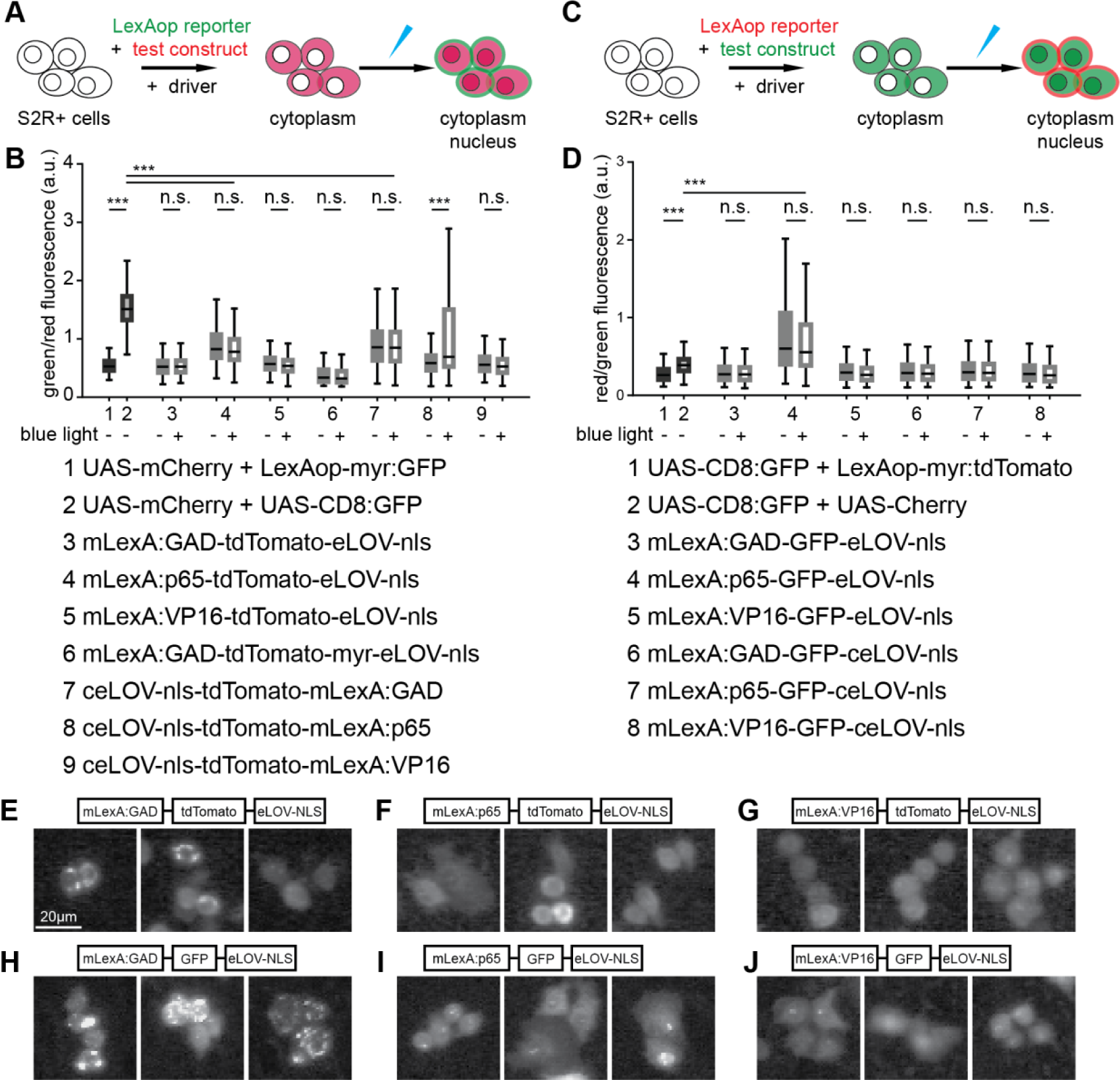
Design of a light-gated expression system based on LOV (suppl. to Figure 1). **A.** S2R+ cells were transfected with the driver pMET-GAL4, the LexAop reporter LexAop-myr:GFP and the test construct. Cells were then kept in the dark or exposed to light. **B.** Ratio of LexAop reporter myr:GFP expression in relation to expression of the different test constructs determined by tdTomato fluorescence. As in Figure 1B, co-transfection of UAS-mCherry with the LexAop reporter LexAop-myr:GFP established the baseline (first bar in the boxplot), whereas co- transfection of UAS-mCherry and UAS-CD8:GFP was used as an approximate measure of co- expression (second bar in the boxplot). Only construct 8, ceLOV-nls-tdTomato-mLexA:p65 (codon optimized ceLOV), elicits expression of the reporter upon light exposure. The tests constructs are indicated below the boxplot; ceLOV stands for eLOV codon optimized for *Drosophila*. At least 200 cells with medium levels of expression of mCherry or tdTomato, from at 2 to 5 transfections of S2R+ cells are represented for each condition. *** represents p values < 0.001, n.s. represents p values > 0.05, obtained with Student’s t test. **C.** S2R+ cells were transfected with a red reporter, LexAop-myr:Tomato, together with test constructs bearing GFP as a tag for visualization. **D.** Ratio of LexAop reporter myr:tdTomato expression in relation to expression of test constructs tagged with GFP. As above, co-transfection of UAS-mCherry and UAS-CD8:GFP served as an approximate measure of co-expression. Co-transfection of UAS-CD8:GFP with the LexAop-myr:tdTomato indicated baseline expression levels. At least 200 cells with medium levels of expression of mCherry or tdTomato, from at 2 to 5 transfections of S2R+ cells are represented for each condition. *** represents p values < 0.001, n.s. represents p values > 0.05, obtained with Student’s t test. **E-J.** Representative examples of S2R+ cells expressing mLexA:GAD-tdTomato- eLOV-nls (E), mLexA:p65-tdTomato-eLOV-nls (F), mLexA:VP16-tdTomato-eLOV-nls (G), mLexA:GAD-GFP-eLOV-nls (H), mLexA:p65-GFP-eLOV-nls (I), mLexA:VP16-GFP-eLOV-nls (J) showing subcellular distribution of each of these combinations. All combinations bearing LexA:GAD form clusters in the cytoplasm, whereas combinations bearing LexA:p65 or LexA:VP16 are more likely to be distributed evenly in the cytoplasm, and sometimes the nucleoplasm.

**Figure S2:**
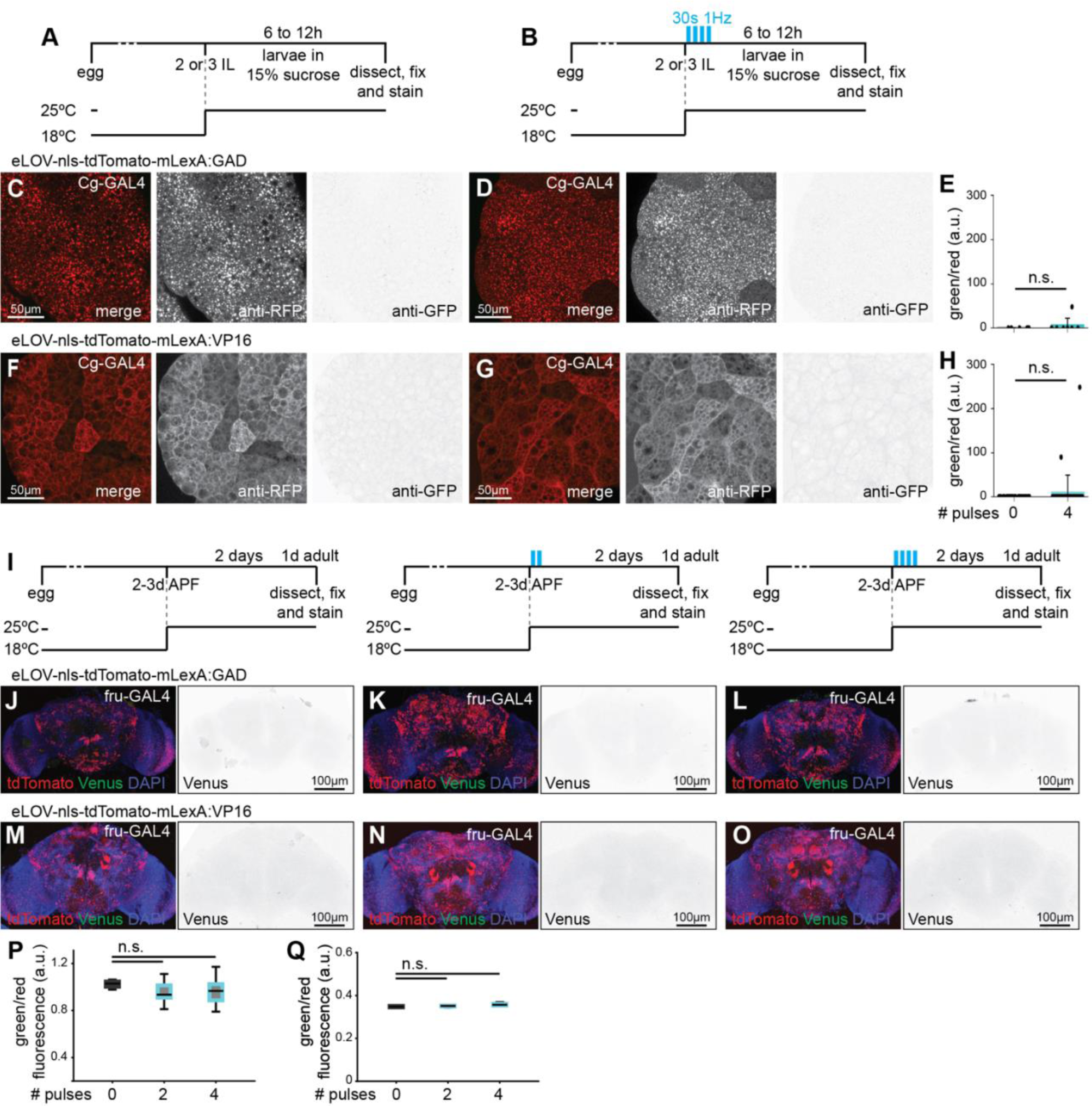
LexA chimeras LexA:GAD and LexA:VP16 combined with N-terminal eLOV are unable to elicit expression of LexAop reporter (suppl. To Figure 2). **A,B.** Schematic representing the timeline of fly rearing temperature and light delivery for the fat bodies in C to H. **C-H.** Fat bodies of second to third instar larvae expressing eLOV-nls-tdTomato- mLexA:GAD (*w, LexAop-CsChrimson:Venus; Cg-GAL4/+; UAS-eLOV-nls-tdTomato- mLexA:GAD/+* C-E) or eLOV-nls-tdTomato-mLexA:VP16 (*w, LexAop-CsChrimson:Venus; Cg-GAL4/+; UAS-eLOV-nls-tdTomato-mLexA:VP16/+* F-H), kept in the dark (C, F) or exposed to four pulses of blue light (each pulse lasting 30s at 1Hz), and incubated 12h at 25°C. The ratio of pixel intensity of anti-GFP signal (LexAop reporter)/anti-RFP signal (test construct) for stained fat bodies is shown for eLOV-nls-tdTomato-mLexA:GAD in E (dark N=3, light N=6) and eLOV- nls-tdTomato-mLexA:VP16 in H (dark N=3, light N=4); n.s. represents p value > 0.05, obtained with Student’s t test. **I.** Schematic representing the timeline of fly rearing temperature and light delivery for the brains in J to Q. **J-Q.** Adult brains showing native expression of LOV-LexA (red) and LexAop-CsChrimson:Venus (Venus, green in merge, and isolated dedicated image on the right) from *w, LexAop-CsChrimson:Venus;+;UAS--eLOV-nls-tdTomato-mLexA:GAD/fru-GAL4* (J-L, P) and *w, LexAop-CsChrimson:Venus;+;UAS--eLOV-nls-tdTomato-mLexA:VP16/fru-GAL4* (M-O, Q) pupae kept in the dark (J, M) or exposed to light (K,L, N, O) at 3-4 days APF, as shown in I. **P,Q.** Ratio of native green (Venus) and red (tdTomato in test construct) fluorescence for cell bodies. Different protocols for light delivery failed to elicit Venus expression in brains expressing eLOV-nls-tdTomato-mLexA:GAD (dark N=3, 2 light pulses N=10, 4 light pulses N=7) or eLOV- nls-tdTomato-mLexA:VP16 (dark N=2, 2 light pulses N=5, 4 light pulses N=4) under control of *fru-GAL4*; n.s. represents p value > 0.05, obtained with Student’s t test.

**Figure S3:**
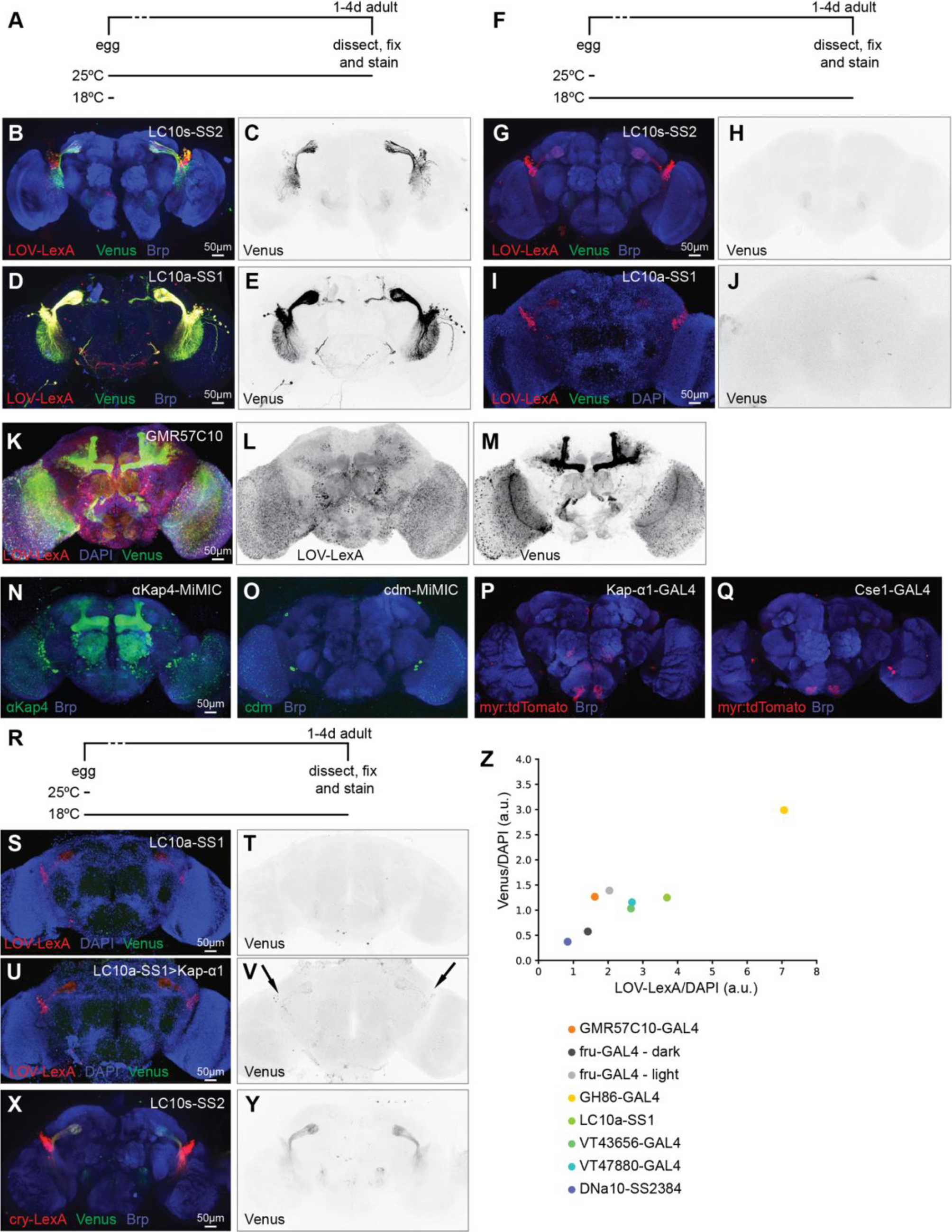
LOV-LexA tests in neurons. **A.** Schematic representing the timeline of fly rearing for the brains shown in B to E. **B-E.** LOV- LexA under control of *LC10s-SS2* (B, N=4) or *LC10a-SS1* (D, N=4) driver in flies reared at 25°C shows expression of the LexAop-reporter transgene Venus (C, E). **F.** Schematic representing the timeline of fly rearing for the brains shown in G to J. **G-J.** LOV-LexA under control of *LC10s- SS2* (G, N=5) or *LC10a-SS1* (I, N=2) driver in flies reared at 18°C shows no expression of the LexAop-reporter transgene Venus (H, J). **K-M.** LOV-LexA (K, L) under control of a panneuronal driver GMR57C10-GAL4 showing LOV-LexA distribution predominantly in cell bodies (L) and uncorrelated expression of the LexAop-reporter Venus (M) in different neurons, N=6. **N, O.** Expression of the EGFP-tagged importins *ɑKap4* and *cdm* with MiMIC lines *ɑKap4^MI0631^*(N=6), and *cdm^MI06239^* (N=4). **P, Q.** Expression pattern of importins *Kap-ɑ1* and *Cse1* visualized with myr:tdTomato under control of GAL4 inserted into *Kap-ɑ1* (N=4) and *Cse1* (N=5) gene loci. **R.** Schematic representing the timeline of fly rearing for the brains shown in S to Y. **S-V.** Ectopic expression of *Kap-ɑ1* in flies reared at 18°C and kept in the dark renders LOV-LexA leaky (U, V with N=3) compared to expression of LOV-LexA alone under the same conditions (S, T, with N=5). **X-Y.** Flies expressing cryptochrome split-LexA reared at 18°C in the dark present expression of the LexAop reporter Venus. **Z.** Average ratio of Venus native signal intensity relative to DAPI signal intensity in relation to the average ratio of tdTomato native signal intensity relative to DAPI signal intensity for flies raised at 18°C and kept in the dark (except for fru-GAL4 – light), to test the dark state of LOV-LexA for various drivers of different strength shows that above certain levels of expression, Venus expression correlates with LOV-LexA expression level, even in the absence of light exposure. If expressed at moderate to low levels, LOV-LexA maintains low transcriptional activity, that increases with light exposure, as is the case for *fru-GAL4* (N=12 exposed to light, N=13 kept in the dark, see Figure 3 D-H). For other drivers, N=2 to 5 brains from flies raised in the dark at 18°C, are plotted.

**Table S1.** Genetic constructs used in this study.

| Name | Features | Vector | Promoter | Details |
| --- | --- | --- | --- | --- |
| c204 | tdTomato-mLexA:GAD | pJFRC7 | UAS | modified LexA:GAD codon optimized for *Drosophila* |
| c205 | tdTomato-LexA:GAD | pJFRC7 | UAS | LexA:GAD codon optimized for *Drosophila* |
| c214 | tdTomato-mLexA:p65 | pJFRC7 | UAS | Modified LexA:p65 codon optimized for *Drosophila* |
| c215 | tdTomato-LexA:p65 | pJFRC7 | UAS | LexA:p65 codon optimized for *Drosophila* |
| c224 | tdTomato-mLexA:VP16 | pJFRC7 | UAS | Modified LexA:VP16 codon optimized for *Drosophila* |
| c225 | tdTomato-LexA:VP16 | pJFRC7 | UAS | LexA:VP16 codon optimized for *Drosophila* |
| c11 | eLOV-nls-tdTomato-mLexA:GAD | pJFRC7 | UAS | LexA:GAD codon optimized for *Drosophila*, with NLS-like modified |
| c12 | eLOV-nls-tdTomato-mLexA:p65 | pJFRC7 | UAS | LexA:p65 codon optimized for *Drosophila*, with NLS-like modified |
| c13 | eLOV-nls-tdTomato-mLexA:VP16 | pJFRC7 | UAS | LexA:VP16 codon optimized for *Drosophila*, with NLS-like modified |
| c17 | mLexA:GAD-tdTomato-eLOV-nls | pJFRC7 | UAS | LexA:GAD codon optimized for *Drosophila*, with NLS-like modified |
| c18 | mLexA:p65-tdTomato-eLOV-nls | pJFRC7 | UAS | LexA:p65 codon optimized for *Drosophila*, with NLS-like modified |
| c19 | mLexA:VP16-tdTomato-eLOV-nls | pJFRC7 | UAS | LexA:VP16 codon optimized for *Drosophila*, with NLS-like modified |
| c111 | ceLOV-nls-tdTomato-mLexA:GAD | pJFRC7 | UAS | LexA:GAD codon optimized for *Drosophila*, with NLS-like modified<br>eLOV evolved LOV codon optimized for *Drosophila* |
| c121 | ceLOV-nls-tdTomato-mLexA:p65 | pJFRC7 | UAS | LexA:p65 codon optimized for *Drosophila*, with NLS-like modified<br>eLOV evolved LOV codon optimized for *Drosophila* |
| c131 | ceLOV-nls-tdTomato-mLexA:VP16 | pJFRC7 | UAS | LexA:VP16 codon optimized for *Drosophila*, with NLS-like modified<br>eLOV evolved LOV codon optimized for *Drosophila* |
| c26 | mLexA:GAD-GFP-eLOV-nls | pJFRC7 | UAS | LexA:GAD codon optimized for *Drosophila*, with NLS-like modified |
| c27 | mLexA:p65-GFP-eLOV-nls | pJFRC7 | UAS | LexA:p65 codon optimized for *Drosophila*, with NLS-like modified |
| c28 | mLexA:VP16-GFP-eLOV-nls | pJFRC7 | UAS | LexA:VP16 codon optimized for *Drosophila*, with NLS-like modified |
| c29 | mLexA:GAD-GFP-ceLOV-nls | pJFRC7 | UAS | LexA:GAD codon optimized for *Drosophila*, with NLS-like modified<br>eLOV evolved LOV codon optimized for *Drosophila* |
| c30 | mLexA:p65-GFP-ceLOV-nls | pJFRC7 | UAS | LexA:p65 codon optimized for *Drosophila*, with NLS-like modified<br>eLOV evolved LOV codon optimized for *Drosophila* |
| c31 | mLexA:VP16-GFP-ceLOV-nls | pJFRC7 | UAS | LexA:VP16 codon optimized for *Drosophila*, with NLS-like modified<br>eLOV evolved LOV codon optimized for *Drosophila* |
| myrTom | LexAop-myr:tdTomato | pJFRC19 | LexAop | LexAop driving tdTomato expression; reporter for activity of LexA-transactivator-eLOV-tag with GFP or FLAG |
| mCherry | mCherry | pJFRC7 | UAS | UAS driving mCherry expression; reporter for transfection efficiency and negative control for LexA-transactivator-eLOV-tag constructs |
| myrGFP | LexAop-myr:GFP | pJFRC19 | LexAop | Pfeiffer, et al, 2008 and 2010<br>Addgene # 26224 |
| CD8:GFP | CD8:GFP | pJFRC7 | UAS | Pfeiffer, et al, 2008 and 2010<br>Addgene # 26220 |

**Table S2.** Drosophila melanogaster strains used in this study.

| *Drosophila melanogaster* strains | Reference | Source |
| --- | --- | --- |
| *Cg-GAL4* | Asha, et al., 2003 | Bloomington Drosophila Stock Center 7011 |
| *fru-GAL4* | Stockinger, et al., 2005 | Barry J. Dickson |
| *GH86-GAL4* | Bloomington Drosophila Stock Center | Bloomington Drosophila Stock Center 36339 |
| *vGlut-GAL4* | Bloomington Drosophila Stock Center | Bloomington Drosophila Stock Center |
| *GMR57C10-GAL4* | Jennet, et al., 2012 | FlyLight, Janelia Research Campus |
| *SS00324* | Jennet, et al., 2012 | FlyLight, Janelia Research Campus |
| *LC10a-SS1* | Ribeiro, et al., 2018 | VDRC |
| *VT043656-GAL4* | Tirian and Dickson, 2017 | VDRC |
| *VT047880-GAL4* | Tirian and Dickson, 2017 | VDRC |
| *LC10s-SS2* | Ribeiro, et al., 2018 | VDRC |
| *UAS-LOV-LexA* in attP1 su(Hw) | this study | VDRC, stock number 311200 |
| *UAS-eLOV-nls-tdTomato-mLexA:GAD* in attP1 su(Hw) | this study | - |
| *UAS-eLOV-nls-tdTomato-mLexA:VP16* in attP1 su(Hw) | this study | - |
| *LexAop-CsChrimson:Venus* in attP18 | Klapoetke, et al., 2014 | Vivek Jayaraman, Janelia Research Campus |
| *LexAop-myr:GFP* | Pfeifer, et al., 2010 | Gerald Rubin, Janelia Research Campus |
| *y<sup>1</sup>,w<sup>\*</sup>;Mi[MIC]aKap4<sup>M106313</sup>PVRAP<sup>M106313</sup>* | Venken, et al., 2011 | Bloomington Drosophila Stock Center 41517 |
| *w<sup>\*</sup>; P[UAS- Kap-*al.M*]2* | Wharton, K. (2008.08.22) personal communication to FlyBase | Bloomington Drosophila Stock Center 25399 |
| *w<sup>1118</sup>;Pbac[IT.GAL4]Kap-<sup>a14018-G4</sup>/TM6B, Tb<sup>1</sup>* | Gohl et al., 2011 | Bloomington Drosophila Stock Center 77639 |
| $y^l w^*$; *Mi[MIC]wrd*<sup>MI06239</sup><br>*cdm*<sup>MI06239</sup> | Nagarkar-Jaiswal, et al., 2015 | Bloomington Drosophila Stock Center 59644 |
| $w^{1118}$; *Mi[ET1]Cse1*<sup>MB08748</sup><br>*mdy*<sup>MB08748</sup>/*SM6a* | Bellen, et al., 2011 | Bloomington Drosophila Stock Center 26129 |
| $y^l w^*$;<br>*P[w[+mW.hs]=GawB]109(2)80*<br>, *P[w[+mC]=UAS-*<br>*mCD8::GFP.L]LL5* | Lee and Luo, 1999 | Bloomington Drosophila Stock Center 8768 |

**Table S3.** Protocols for blue light delivery used in this study.

|  | Source of blue light | wavelength | intensity | pulse | Nr of pulses | figures |
| --- | --- | --- | --- | --- | --- | --- |
| S2R+ cells | LED | 460-500nm | 11.7 mW | 1Hz for 30s | 4 to 6, 1 hour apart | 1, S1 |
| Larvae (fat body) | LED | 460-500nm | 11.7 mW | 1Hz for 30s | 3 to 6, 30 min apart | 2, S2 |
| Pupae (oenocytes) | 1-photon | 485nm | 1.62 μW | 0.33Hz for 90s | 1 | 2 |
| Pupae (neurons) | 1-photon | 458nm | 2.38 μW | 0.56Hz for 90s | 4, 10 min apart | 3, S2, S3 |
| Adults (neurons) | 1-photon | 485nm | 2.49 μW | 0.33Hz for 90s | 4, 10 to 20 min apart | 4 |

## References

Asha, H., I. Nagy, G. Kovacs, D. Stetson, I. Ando et al., 2003 Analysis of Ras-induced overproliferation in Drosophila hemocytes. Genetics 163: 203–215.

Bath, D. E., J. R. Stowers, D. Hormann, A. Poehlmann, B. J. Dickson et al., 2014 FlyMAD: rapid thermogenetic control of neuronal activity in freely walking Drosophila. Nat Methods 11: 756–762.

Bellen, H. J., C. J. O’Kane, C. Wilson, U. Grossniklaus, R. K. Pearson et al., 1989 P-element-mediated enhancer detection: a versatile method to study development in Drosophila. Genes Dev 3: 1288–1300.

Brand, A. H., and N. Perrimon, 1993 Targeted gene expression as a means of altering cell fates and generating dominant phenotypes. Development 118: 401–415.

Cachero, S., A. D. Ostrovsky, J. Y. Yu, B. J. Dickson and G. S. Jefferis, 2010 Sexual dimorphism in the fly brain. Curr Biol 20: 1589–1601.

Carpenter, A. E., T. R. Jones, M. R. Lamprecht, C. Clarke, I. H. Kang et al., 2006 CellProfiler: image analysis software for identifying and quantifying cell phenotypes. Genome Biol 7: R100.

Cavanaugh, K. E., M. F. Staddon, E. Munro, S. Banerjee and M. L. Gardel, 2020 RhoA mediates epithelial cell shape changes via mechanosensitive endocytosis. Dev Cell 52: 152–166 e155.

Chan, Y. B., O. V. Alekseyenko and E. A. Kravitz, 2015 Optogenetic control of gene expression in Drosophila. PLoS One 10: e0138181.

Chen, I. W., E. Papagiakoumou and V. Emiliani, 2018 Towards circuit optogenetics. Curr Opin Neurobiol 50: 179–189.

Christie, J. M., P. Reymond, G. K. Powell, P. Bernasconi, A. A. Raibekas et al., 1998 Arabidopsis NPH1: a flavoprotein with the properties of a photoreceptor for phototropism. Science 282: 1698–1701.

Christie, J. M., M. Salomon, K. Nozue, M. Wada and W. R. Briggs, 1999 LOV (light, oxygen, or voltage) domains of the blue-light photoreceptor phototropin (nph1): binding sites for the chromophore flavin mononucleotide. Proc Natl Acad Sci U S A 96: 8779–8783.

Costa, M., J. D. Manton, A. D. Ostrovsky, S. Prohaska and G. S. Jefferis, 2016 NBLAST: rapid, sensitive comparison of neuronal structure and construction of neuron family databases. Neuron 91: 293–311.

Crosson, S., and K. Moffat, 2002 Photoexcited structure of a plant photoreceptor domain reveals a light-driven molecular switch. Plant Cell 14: 1067–1075.

de Mena, L., and D. E. Rincon-Limas, 2020 PhotoGal4: A versatile light-dependent switch for spatiotemporal control of gene expression in Drosophila explants. iScience 23: 101308.

de Mena, L., P. Rizk and D. E. Rincon-Limas, 2018 Bringing light to transcription: the optogenetics repertoire. Front Genet 9: 518.

del Valle Rodriguez, A. D. Didiano and C. Desplan, 2011 Power tools for gene expression and clonal analysis in Drosophila. Nat Methods 9: 47–55.

di Pietro, F., S. Herszterg, A. Huang, F. Bosveld, C. Alexandre et al., 2021 Rapid and robust optogenetic control of gene expression in Drosophila. Dev Cell 56: 3393–3404 e3397.

Di Ventura, B., and B. Kuhlman, 2016 Go in! Go out! Inducible control of nuclear localization. Curr Opin Chem Biol 34: 62–71.

Diensthuber, R. P., C. Engelhard, N. Lemke, T. Gleichmann, R. Ohlendorf et al., 2014 Biophysical, mutational, and functional investigation of the chromophore-binding pocket of light-oxygen-voltage photoreceptors. ACS Synth Biol 3: 811–819.

Dionne, H., K. L. Hibbard, A. Cavallaro, J. C. Kao and G. M. Rubin, 2018 Genetic reagents for making split-GAL4 lines in Drosophila. Genetics 209: 31–35.

Dolan, M. J., S. Frechter, A. S. Bates, C. Dan, P. Huoviala et al., 2019 Neurogenetic dissection of the Drosophila lateral horn reveals major outputs, diverse behavioural functions, and interactions with the mushroom body. Elife 8.

Echalier, G., 1997 Drosophila Cells in Culture. Academic Press, New York. 702 pp.

Emelyanov, A., and S. Parinov, 2008 Mifepristone-inducible LexPR system to drive and control gene expression in transgenic zebrafish. Dev Biol 320: 113–121.

Feng, K., M. T. Palfreyman, M. Hasemeyer, A. Talsma and B. J. Dickson, 2014 Ascending SAG neurons control sexual receptivity of Drosophila females. Neuron 83: 135–148.

Feng, K., R. Sen, R. Minegishi, M. Dubbert, T. Bockemuhl et al., 2020 Distributed control of motor circuits for backward walking in Drosophila. Nat Commun 11: 6166.

Fischbach, K.-F., and A. P. M. Dittrich, 1989 The optic lobe of *Drosophila melanogaster*. I. A Golgi analysis of wild-type structure. Cell Tissue Res 258: 441–475.

Gailey, D. A., and J. C. Hall, 1989 Behavior and cytogenetics of fruitless in Drosophila melanogaster: different courtship defects caused by separate, closely linked lesions. Genetics 121: 773–785.

Gibson, D. G., L. Young, R. Y. Chuang, J. C. Venter, C. A. Hutchison, 3rd et al., 2009 Enzymatic assembly of DNA molecules up to several hundred kilobases. Nat Methods 6: 343–345.

Golic, K. G., and S. Lindquist, 1989 The FLP recombinase of yeast catalyzes site-specific recombination in the Drosophila genome. Cell 59: 499–509.

Grossniklaus, U., H. J. Bellen, C. Wilson and W. J. Gehring, 1989 P-element-mediated enhancer detection applied to the study of oogenesis in Drosophila. Development 107: 189–200.

Guntas, G., R. A. Hallett, S. P. Zimmerman, T. Williams, H. Yumerefendi et al., 2015 Engineering an improved light-induced dimer (iLID) for controlling the localization and activity of signaling proteins. Proc Natl Acad Sci U S A 112: 112–117.

Hadjieconomou, D., S. Rotkopf, C. Alexandre, D. M. Bell, B. J. Dickson et al., 2011 Flybow: genetic multicolor cell labeling for neural circuit analysis in Drosophila melanogaster. Nat Methods 8: 260–266.

Harper, S. M., L. C. Neil and K. H. Gardner, 2003 Structural basis of a phototropin light switch. Science 301: 1541–1544.

Hindmarsh Sten, T. R. Li, A. Otopalik and V. Ruta, 2021 Sexual arousal gates visual processing during Drosophila courtship. Nature 595: 549–553.

Horii, T., T. Ogawa and H. Ogawa, 1981 Nucleotide sequence of the lexA gene of E. coli. Cell 23: 689–697.

Hosokawa, K., M. Nishi, H. Sakamoto, Y. Tanaka and M. Kawata, 2008 Regional distribution of importin subtype mRNA expression in the nervous system: study of early postnatal and adult mouse. Neuroscience 157: 864–877.

Huala, E., P. W. Oeller, E. Liscum, I. S. Han, E. Larsen et al., 1997 Arabidopsis NPH1: a protein kinase with a putative redox-sensing domain. Science 278: 2120–2123.

Huberman, A. D., M. B. Feller and B. Chapman, 2008 Mechanisms underlying development of visual maps and receptive fields. Annu Rev Neurosci 31: 479–509.

Inagaki, H. K., Y. Jung, E. D. Hoopfer, A. M. Wong, N. Mishra et al., 2014 Optogenetic control of Drosophila using a red-shifted channelrhodopsin reveals experience-dependent influences on courtship. Nat Methods 11: 325–332.

Isaacman-Beck, J., K. C. Paik, C. F. R. Wienecke, H. H. Yang, Y. E. Fisher et al., 2020 SPARC enables genetic manipulation of precise proportions of cells. Nat Neurosci 23: 1168–1175.

Jang, A. R. K. Moravcevic, L. Saez, M. W. Young and A. Sehgal, 2015 Drosophila TIM binds importin alpha1, and acts as an adapter to transport PER to the nucleus. PLoS Genet 11: e1004974.

Jayaraman, P., K. Devarajan, T. K. Chua, H. Zhang, E. Gunawan et al., 2016 Blue light-mediated transcriptional activation and repression of gene expression in bacteria. Nucleic Acids Res 44: 6994–7005.

Jenett, A., G. M. Rubin, T. T. Ngo, D. Shepherd, C. Murphy et al., 2012 A GAL4-driver line resource for Drosophila neurobiology. Cell Rep 2: 991–1001.

Kawano, F., H. Suzuki, A. Furuya and M. Sato, 2015 Engineered pairs of distinct photoswitches for optogenetic control of cellular proteins. Nat Commun 6: 6256.

Kim, C. K. K. F. Cho, M. W. Kim and A. Y. Ting, 2019 Luciferase-LOV BRET enables versatile and specific transcriptional readout of cellular protein-protein interactions. Elife 8.

Kim, M. W. W. Wang, M. I. Sanchez, R. Coukos, M. von Zastrow et al., 2017a Time-gated detection of protein-protein interactions with transcriptional readout. Elife 6.

Kim, S. S. H. Rouault, S. Druckmann and V. Jayaraman, 2017b Ring attractor dynamics in the Drosophila central brain. Science 356: 849–853.

Klapoetke, N. C., Y. Murata, S. S. Kim, S. R. Pulver, A. Birdsey-Benson et al., 2014 Independent optical excitation of distinct neural populations. Nat Methods 11: 338–346.

Kockel, L., C. Griffin, Y. Ahmed, L. Fidelak, A. Rajan et al., 2019 An interscholastic network to generate LexA enhancer trap lines in Drosophila. G3 (Bethesda) 9: 2097–2106.

Kockel, L., L. M. Huq, A. Ayyar, E. Herold, E. MacAlpine et al., 2016 A Drosophila LexA enhancer-trap resource for developmental biology and neuroendocrine research. G3 (Bethesda) 6: 3017–3026.

Kvon, E. Z., T. Kazmar, G. Stampfel, J. O. Yanez-Cuna, M. Pagani et al., 2014 Genome-scale functional characterization of Drosophila developmental enhancers in vivo. Nature 512: 91–95.

Lai, S. L., and T. Lee, 2006 Genetic mosaic with dual binary transcriptional systems in Drosophila. Nat Neurosci 9: 703–709.

Larkin, A., S. J. Marygold, G. Antonazzo, H. Attrill, G. Dos Santos et al., 2021 FlyBase: updates to the Drosophila melanogaster knowledge base. Nucleic Acids Res 49: D899–D907.

Lee, T., and L. Luo, 1999 Mosaic analysis with a repressible cell marker for studies of gene function in neuronal morphogenesis. Neuron 22: 451–461.

Lee, T., and L. Luo, 2001 Mosaic analysis with a repressible cell marker (MARCM) for Drosophila neural development. Trends Neurosci 24: 251–254.

Lindquist, S., 1986 The heat-shock response. Annu Rev Biochem 55: 1151–1191.

Lis, J. T., J. A. Simon and C. A. Sutton, 1983 New heat shock puffs and beta-galactosidase activity resulting from transformation of Drosophila with an hsp70-lacZ hybrid gene. Cell 35: 403–410.

Loewer, A., P. Soba, K. Beyreuther, R. Paro and G. Merdes, 2004 Cell-type-specific processing of the amyloid precursor protein by Presenilin during Drosophila development. EMBO Rep 5: 405–411.

Lu, B., A. LaMora, Y. Sun, M. J. Welsh and Y. Ben-Shahar, 2012 ppk23-Dependent chemosensory functions contribute to courtship behavior in Drosophila melanogaster. PLoS Genet 8: e1002587.

Luan, H., N. C. Peabody, C. R. Vinson and B. H. White, 2006 Refined spatial manipulation of neuronal function by combinatorial restriction of transgene expression. Neuron 52: 425–436.

Lungu, O. I., R. A. Hallett, E. J. Choi, M. J. Aiken, K. M. Hahn et al., 2012 Designing photoswitchable peptides using the AsLOV2 domain. Chem Biol 19: 507–517.

Makki, R., E. Cinnamon and A. P. Gould, 2014 The development and functions of oenocytes. Annu Rev Entomol 59: 405–425.

Masuyama, K., Y. Zhang, Y. Rao and J. W. Wang, 2012 Mapping neural circuits with activity-dependent nuclear import of a transcription factor. J Neurogenet 26: 89–102.

McGuire, S. E., P. T. Le, A. J. Osborn, K. Matsumoto and R. L. Davis, 2003 Spatiotemporal rescue of memory dysfunction in Drosophila. Science 302: 1765–1768.

McGuire, S. E., G. Roman and R. L. Davis, 2004 Gene expression systems in Drosophila: a synthesis of time and space. Trends Genet 20: 384–391.

McKellar, C. E., J. L. Lillvis, D. E. Bath, J. E. Fitzgerald, J. G. Cannon et al., 2019 Threshold-based ordering of sequential actions during Drosophila courtship. Curr Biol 29: 426–434 e426.

McQuin, C., A. Goodman, V. Chernyshev, L. Kamentsky, B. A. Cimini et al., 2018 CellProfiler 3.0: Next-generation image processing for biology. PLoS Biol 16: e2005970.

Montell, C., 2012 Drosophila visual transduction. Trends Neurosci 35: 356–363.

Motta-Mena, L. B., A. Reade, M. J. Mallory, S. Glantz, O. D. Weiner et al., 2014 An optogenetic gene expression system with rapid activation and deactivation kinetics. Nat Chem Biol 10: 196–202.

Namiki, S., M. H. Dickinson, A. M. Wong, W. Korff and G. M. Card, 2018 The functional organization of descending sensory-motor pathways in Drosophila. Elife 7.

Nassi, J. J., and E. M. Callaway, 2009 Parallel processing strategies of the primate visual system. Nat Rev Neurosci 10: 360–372.

Nern, A., B. D. Pfeiffer and G. M. Rubin, 2015 Optimized tools for multicolor stochastic labeling reveal diverse stereotyped cell arrangements in the fly visual system. Proc Natl Acad Sci U S A 112: E2967–2976.

Niopek, D., D. Benzinger, J. Roensch, T. Draebing, P. Wehler et al., 2014 Engineering light-inducible nuclear localization signals for precise spatiotemporal control of protein dynamics in living cells. Nat Commun 5: 4404.

Niopek, D., P. Wehler, J. Roensch, R. Eils and B. Di Ventura, 2016 Optogenetic control of nuclear protein export. Nat Commun 7: 10624.

Nogi, Y., K. Matsumoto, A. Toh-e and Y. Oshima, 1977 Interaction of super-repressible and dominant constitutive mutations for the synthesis of galactose pathway enzymes in Saccharomyces cerevisiae. Mol Gen Genet 152: 137–144.

Nonet, M. L., 2020 Efficient transgenesis in Caenorhabditis elegans using Flp recombinase-mediated cassette exchange. Genetics 215: 903–921.

Otsuna, H., and K. Ito, 2006 Systematic analysis of the visual projection neurons of Drosophila melanogaster. I. Lobula-specific pathways. J Comp Neurol 497: 928–958.

Panser, K., L. Tirian, F. Schulze, S. Villalba, G. Jefferis et al., 2016 Automatic segmentation of Drosophila neural compartments using GAL4 expression data reveals novel visual pathways. Curr Biol 26: 1943–1954.

Perrimon, N., E. Noll, K. McCall and A. Brand, 1991 Generating lineage-specific markers to study Drosophila development. Dev Genet 12: 238–252.

Petersen, C. C. H., 2019 Sensorimotor processing in the rodent barrel cortex. Nat Rev Neurosci 20: 533–546.

Pfeiffer, B. D., A. Jenett, A. S. Hammonds, T. T. Ngo, S. Misra et al., 2008 Tools for neuroanatomy and neurogenetics in Drosophila. Proc Natl Acad Sci U S A 105: 9715–9720.

Pfeiffer, B. D., T. T. Ngo, K. L. Hibbard, C. Murphy, A. Jenett et al., 2010 Refinement of tools for targeted gene expression in Drosophila. Genetics 186: 735–755.

Potter, C. J., B. Tasic, E. V. Russler, L. Liang and L. Luo, 2010 The Q system: a repressible binary system for transgene expression, lineage tracing, and mosaic analysis. Cell 141: 536–548.

Reade, A., L. B. Motta-Mena, K. H. Gardner, D. Y. Stainier, O. D. Weiner et al., 2017 TAEL: a zebrafish-optimized optogenetic gene expression system with fine spatial and temporal control. Development 144: 345–355.

Rhee, Y., F. Gurel, Y. Gafni, C. Dingwall and V. Citovsky, 2000 A genetic system for detection of protein nuclear import and export. Nat Biotechnol 18: 433–437.

Riabinina, O., D. Luginbuhl, E. Marr, S. Liu, M. N. Wu et al., 2015 Improved and expanded Q-system reagents for genetic manipulations. Nat Methods 12: 219–222, 215 p following 222.

Riabinina, O., and C. J. Potter, 2016 The Q-System: A versatile expression system for Drosophila. Methods Mol Biol 1478: 53–78.

Ribeiro, I. M. A., M. Drews, A. Bahl, C. Machacek, A. Borst et al., 2018 Visual projection neurons mediating directed courtship in Drosophila. Cell 174: 607–621 e618.

Robie, A. A., J. Hirokawa, A. W. Edwards, L. A. Umayam, A. Lee et al., 2017 Mapping the neural substrates of behavior. Cell 170: 393–406 e328.

Rubin, G. M., and A. C. Spradling, 1982 Genetic transformation of Drosophila with transposable element vectors. Science 218: 348–353.

Salinas, F., V. Rojas, V. Delgado, J. Lopez, E. Agosin et al., 2018 Fungal Light-Oxygen-Voltage domains for optogenetic control of gene expression and flocculation in yeast. mBio 9.

Salomon, M., J. M. Christie, E. Knieb, U. Lempert and W. R. Briggs, 2000 Photochemical and mutational analysis of the FMN-binding domains of the plant blue light receptor, phototropin. Biochemistry 39: 9401–9410.

Schindelin, J., I. Arganda-Carreras, E. Frise, V. Kaynig, M. Longair et al., 2012 Fiji: an open-source platform for biological-image analysis. Nat Methods 9: 676–682.

Schretter, C. E., Y. Aso, A. A. Robie, M. Dreher, M. J. Dolan et al., 2020 Cell types and neuronal circuitry underlying female aggression in Drosophila. Elife 9.

Sen, R., M. Wu, K. Branson, A. Robie, G. M. Rubin et al., 2017 Moonwalker Descending Neurons Mediate Visually Evoked Retreat in Drosophila. Curr Biol 27: 766–771.

Serebreni, L., and A. Stark, 2021 Insights into gene regulation: From regulatory genomic elements to DNA-protein and protein-protein interactions. Curr Opin Cell Biol 70: 58–66.

Shaner, N. C., R. E. Campbell, P. A. Steinbach, B. N. Giepmans, A. E. Palmer et al., 2004 Improved monomeric red, orange and yellow fluorescent proteins derived from Discosoma sp. red fluorescent protein. Nat Biotechnol 22: 1567–1572.

Smart, A. D., R. A. Pache, N. D. Thomsen, T. Kortemme, G. W. Davis et al., 2017 Engineering a light-activated caspase-3 for precise ablation of neurons in vivo. Proc Natl Acad Sci U S A 114: E8174–E8183.

Spradling, A. C., and G. M. Rubin, 1982 Transposition of cloned P elements into Drosophila germ line chromosomes. Science 218: 341–347.

Sterne, G. R., H. Otsuna, B. J. Dickson and K. Scott, 2021 Classification and genetic targeting of cell types in the primary taste and premotor center of the adult Drosophila brain. Elife 10.

Stockinger, P., D. Kvitsiani, S. Rotkopf, L. Tirian and B. J. Dickson, 2005 Neural circuitry that governs Drosophila male courtship behavior. Cell 121: 795–807.

Strickland, D., Y. Lin, E. Wagner, C. M. Hope, J. Zayner et al., 2012 TULIPs: tunable, light-controlled interacting protein tags for cell biology. Nat Methods 9: 379–384.

Szuts, D., and M. Bienz, 2000 LexA chimeras reveal the function of Drosophila Fos as a context-dependent transcriptional activator. Proc Natl Acad Sci U S A 97: 5351–5356.

Thistle, R., P. Cameron, A. Ghorayshi, L. Dennison and K. Scott, 2012 Contact chemoreceptors mediate male-male repulsion and male-female attraction during Drosophila courtship. Cell 149: 1140–1151.

Tirian, L., and B. J. Dickson, 2017 The VT GAL4, LexA, and split-GAL4 driver line collections for targeted expression in the *Drosophila* nervous system. bioRxiv.

Toda, H., X. Zhao and B. J. Dickson, 2012 The Drosophila female aphrodisiac pheromone activates ppk23(+) sensory neurons to elicit male courtship behavior. Cell Rep 1: 599–607.

van Haren, J., R. A. Charafeddine, A. Ettinger, H. Wang, K. M. Hahn et al., 2018 Local control of intracellular microtubule dynamics by EB1 photodissociation. Nat Cell Biol 20: 252–261.

Velichkova, M., J. Juan, P. Kadandale, S. Jean, I. Ribeiro et al., 2010 Drosophila Mtm and class II PI3K coregulate a PI(3)P pool with cortical and endolysosomal functions. J Cell Biol 190: 407–425.

Venken, K. J., and H. J. Bellen, 2012 Genome-wide manipulations of Drosophila melanogaster with transposons, Flp recombinase, and PhiC31 integrase. Methods Mol Biol 859: 203–228.

Venken, K. J., and H. J. Bellen, 2014 Chemical mutagens, transposons, and transgenes to interrogate gene function in Drosophila melanogaster. Methods 68: 15–28.

Venken, K. J., K. L. Schulze, N. A. Haelterman, H. Pan, Y. He et al., 2011 MiMIC: a highly versatile transposon insertion resource for engineering Drosophila melanogaster genes. Nat Methods 8: 737–743.

Walker, G. C., 1985 Inducible DNA repair systems. Annu Rev Biochem 54: 425–457.

Wang, F., K. Wang, N. Forknall, C. Patrick, T. Yang et al., 2020 Neural circuitry linking mating and egg laying in Drosophila females. Nature 579: 101–105.

Wang, W., C. P. Wildes, T. Pattarabanjird, M. I. Sanchez, G. F. Glober et al., 2017 A light- and calcium-gated transcription factor for imaging and manipulating activated neurons. Nat Biotechnol 35: 864–871.

Wang, X., L. He, Y. I. Wu, K. M. Hahn and D. J. Montell, 2010 Light-mediated activation reveals a key role for Rac in collective guidance of cell movement in vivo. Nat Cell Biol 12: 591–597.

Wilson, C., R. K. Pearson, H. J. Bellen, C. J. O’Kane, U. Grossniklaus et al., 1989 P-element-mediated enhancer detection: an efficient method for isolating and characterizing developmentally regulated genes in Drosophila. Genes Dev 3: 1301–1313.

Wu, M., A. Nern, W. R. Williamson, M. M. Morimoto, M. B. Reiser et al., 2016 Visual projection neurons in the Drosophila lobula link feature detection to distinct behavioral programs. Elife 5.

Xu, T., and G. M. Rubin, 1993 Analysis of genetic mosaics in developing and adult Drosophila tissues. Development 117: 1223–1237.

Yagi, R., F. Mayer and K. Basler, 2010 Refined LexA transactivators and their use in combination with the Drosophila Gal4 system. Proc Natl Acad Sci U S A 107: 16166–16171.

Yamamoto, N., and X. W. Deng, 1999 Protein nucleocytoplasmic transport and its light regulation in plants. Genes Cells 4: 489–500.

Yanez-Cuna, J. O., C. D. Arnold, G. Stampfel, L. M. Boryn, D. Gerlach et al., 2014 Dissection of thousands of cell type-specific enhancers identifies dinucleotide repeat motifs as general enhancer features. Genome Res 24: 1147–1156.

Yu, J. Y., M. I. Kanai, E. Demir, G. S. Jefferis and B. J. Dickson, 2010 Cellular organization of the neural circuit that drives Drosophila courtship behavior. Curr Biol 20: 1602–1614.

Yumerefendi, H., D. J. Dickinson, H. Wang, S. P. Zimmerman, J. E. Bear et al., 2015 Control of protein activity and cell fate specification via light-mediated nuclear translocation. PLoS One 10: e0128443.

Yumerefendi, H., A. M. Lerner, S. P. Zimmerman, K. Hahn, J. E. Bear et al., 2016 Light-induced nuclear export reveals rapid dynamics of epigenetic modifications. Nat Chem Biol 12: 399–401.

Zayner, J. P., C. Antoniou and T. R. Sosnick, 2012 The amino-terminal helix modulates light-activated conformational changes in AsLOV2. J Mol Biol 419: 61–74.

Zhao, E. M., Y. Zhang, J. Mehl, H. Park, M. A. Lalwani et al., 2018 Optogenetic regulation of engineered cellular metabolism for microbial chemical production. Nature 555: 683–687.

